# Intraspecific variation of transposable elements reveals differences in the evolutionary history of fungal phytopathogen pathotypes

**DOI:** 10.1101/2022.11.27.518126

**Authors:** Anne A. Nakamoto, Pierre M. Joubert, Ksenia V. Krasileva

## Abstract

Transposable elements (TEs) contribute to intraspecific variation and play important roles in the evolution of fungal genomes. However, our understanding of the processes that shape TE landscapes is limited, as is our understanding of the relationship between TE content, population structure, and evolutionary history of fungal species. Fungal plant pathogens, which often have host-specific populations, are useful systems in which to study intraspecific TE content diversity. Here, we describe TE dynamics in five lineages of *Magnaporthe oryzae*, the fungus that causes blast disease of rice, wheat, and many other grasses. We identified differences in TE content across these lineages, and showed that recent lineage-specific expansions of certain TEs have contributed to overall greater TE content in rice-infecting and *Setaria*-infecting lineages. We reconstructed the evolutionary histories of LTR-retrotransposon expansions and found that in some cases they were caused by complex proliferation dynamics of one element, and in others by multiple elements from an older population of TEs multiplying in parallel. Additionally, we found evidence suggesting the recent transfer of a DNA transposon between rice and wheat-infecting *M. oryzae* lineages, and a region showing evidence of homologous recombination between those lineages, which could have facilitated such a transfer. By investigating intraspecific TE content variation, we uncovered key differences in the proliferation dynamics of TEs in various pathotypes of a fungal plant pathogen, giving us a better understanding of the evolutionary history of the pathogen itself.

## Introduction

Many fungal species display extensive intraspecific variation, allowing them to adapt to a wide range of lifestyles and environments (Mathieu et al. 2018; Branco 2019; Monte et al. 2021). Transposable elements (TEs), a diverse collection of repetitive, mobile sequences, are known to generate genomic diversity and contribute to genome evolution (Wells and Feschotte 2020; Almojil et al. 2021). Along with the substantial diversity of TE content across fungal species (Raffaele and Kamoun 2012; Castanera et al. 2016; Muszewska et al. 2019), there are examples of intraspecific TE content variation in edible mushrooms, fungal pathogens, mycorrhizal fungi, and yeast (Castanera et al. 2016; Oggenfuss et al. 2021; Gourlie et al. 2022; Shirke et al. 2016; Chen et al. 2018; Bleykasten-Grosshans et al. 2021). Yet, we are only beginning to understand how differences in TE content arise in such systems, and how this may reflect a species’ evolutionary history. Fungal plant pathogens provide interesting models for investigating TE content diversity, as many have host-specific populations (Möller and Stukenbrock 2017). Many of these fungi are also thought to have a “two-speed” genome structure, where slowly evolving, gene-rich regions are separated from rapidly evolving regions with many TEs and few genes (Dong et al. 2015; Faino et al. 2016). Disease-causing effector genes are thought to undergo rapid gain, loss, and evolution in genomic regions associated with TEs, which can benefit the pathogen by allowing evasion of their host’s immune response (Sánchez-Vallet et al. 2018). Although some studies have identified intraspecific differences in fungal plant pathogen TE content (Shirke et al. 2016; Oggenfuss et al. 2021; Gourlie et al. 2022), the relationship between TEs and the evolutionary histories of these species remains largely uncharacterized.

*Magnaporthe oryzae* is an important fungal pathogen that causes the blast disease of various grasses, including important crops such as rice and wheat (Dean et al. 2012: 10). *M. oryzae’s* wide host range is associated with substantial intraspecific diversity. The species is composed of distinct pathotypes that include *Oryza* (rice), *Setaria* (foxtail), *Triticum* (wheat), *Lolium* (ryegrass), and *Eleusine* (goosegrass) - infecting lineages (MoO, MoS, MoT, MoL, and MoE, respectively) (Gladieux, Condon, et al. 2018). All lineages are thought to have arisen recently, yet there is large variation in their relative ages. It is well-accepted that the closely related MoO and MoS lineages diverged from their common ancestor around the time of rice domestication, approximately 9,800 years ago via a host-shift of MoS to rice (Couch et al. 2005; Gladieux, Ravel, et al. 2018; Zhong et al. 2018). However, wheat blast was discovered much more recently in 1985 (Singh et al. 2021), and was thought to have arisen via a host shift of MoL to wheat (Inoue et al. 2017; Ceresini et al. 2019). Alternatively, a recent study suggests that a large admixture event involving recombination between isolates of multiple pathotypes, including MoE and a relative of MoO and MoS, may have given rise to the closely related MoT and MoL within the past 60 years (Rahnama et al. 2021). It is clear that we have yet to fully understand the processes shaping the present structure of the *M. oryzae* lineages.

Previous studies have shown that TEs can have major effects on the *M. oryzae* genome and its host specificity. For example, insertion of the *POT2* DNA transposon into the *AVR-Pib* effector gene of MoO isolates allowed them to evade recognition by the *Pib* gene in rice, overcoming resistance (Li et al. 2022). Additionally, it is hypothesized that TE insertions caused the functional loss of the *PWT3* effector, enabling a host jump of MoL to wheat (Inoue et al. 2017). Many studies have also shown that *M. oryzae* experiences frequent gene gains and losses, often in association with TEs (Shirke et al. 2016; Yoshida et al. 2016; Thierry et al. 2022; Joubert and Krasileva 2023). Finally, extrachromosomal circular DNAs have been shown to confer great adaptive potential (Paulsen et al. 2018), and *M. oryzae* was found to produce a large set of eccDNAs consisting of many LTR-retrotransposon sequences (Joubert and Krasileva 2022). Although TEs are associated with many adaptive processes in *M. oryzae*, the relationship between TE content variation and the population structure and evolutionary history of the species remains unknown.

In this study, we compared TE content and proliferation dynamics across the *M. oryzae* lineages. We assembled an unbiased library of TEs and produced robust annotations, which revealed striking differences in overall TE content between lineages. We observed that recent lineage-specific expansions of LTR-retrotransposons have contributed to greater TE content in MoO and MoS. The histories and dynamics of these expansions were complex. Some were caused by the proliferation of one element, while others consisted of multiple elements from an older population of TEs that proliferated in parallel. Additionally, we found evidence suggesting a recent transfer of a DNA transposon between MoO and MoT, and found a potential region of recombination between those lineages that could have facilitated such a transfer. Together, these results showed complex TE expansion dynamics in *M. oryzae* lineages that shaped *M. oryzae’s* evolutionary history.

## Results

### TE content in *M. oryzae* varies greatly across lineages and isolates

To analyze TE content in *M. oryzae*, we first constructed a library representative of TE diversity in all lineages. Since highly contiguous genome assemblies provide the most complete and accurate view of TE content (Rech et al. 2022), a set of highly contiguous *M. oryzae* genomes was gathered from NCBI GenBank for each lineage (Table S1). We then constructed a pipeline based on previous methods (Muszewska et al. 2019) to annotate TEs. The pipeline (Figure S1) utilized one representative genome from each lineage to perform *de novo* repeat annotation. We then added the RepBase (Bao et al. 2015) library of known TEs in fungi, and filtered the combined library against a list of TE-associated protein domains (Muszewska et al. 2019). This ensured that only elements with the potential to be active were kept. The library was further refined by manual classification of elements using protein domain-based phylogenies (Figure S1). Elements that formed a subclade with a known RepBase TE were classified as being part of the same family. This resulted in the classification of many *de novo* elements as members of known families (*Ty3_MAG1, Ty3_MAG2, Grasshopper, MAG_Ty3, MGRL3, PYRET, MGR583, MoTeR1,* and *POT2*), or as new families part of a known TE superfamily (*Copia_elem* and *TcMar_elem*) (Table S2). TEs that did not group in a subclade containing a known family or superfamily were classified as ‘unknown’. We then used the classified library to annotate TEs in our set of *M. oryzae* genomes (Table S1), and verified that each hit contained a TE-associated domain. This approach provided high-quality and unbiased copy number and positional information of TEs in each genome.

Using our TE annotations, we observed striking differences in TE content between genomes of different lineages, and these differences seemed to follow the evolutionary relationships between lineages (Figure 1, Figure S2). Most notably, MoO and MoS genomes contained much higher TE content than MoT, MoL, and MoE (Figure 1A). In MoO and MoS, an average of 11.14% (5.1 Mb) of the genome consisted of annotated TEs, while the average was 5.44% (2.4 Mb) for the other three lineages (Table S1, Figure 1A). We observed that principal component analysis (PCA) of TE content also clearly separates the lineages (Figure S3A-C). The *Magnaporthe grisea* isolate that was used as an outgroup had very similar TE content to the MoT, MoL, and MoE isolates, suggesting that the MoO-MoS clade may have acquired its higher TE content after diverging from the other lineages (Figure 1A). An analysis of the genome sizes of the isolates showed that the increased TE content in MoO-MoS was not due to genome duplication (Table S1, Figure S3D). Although we did observe a correlation between TE content and genome size, most of the signal appears to come from MoO and MoS isolates (Figure S3D). Furthermore, correlation tests showed that the differences we observed between lineages were not due to assembly quality or completeness, while TE content was highly correlated with the lineage an isolate belonged to (Figure S3E). Finally, we observed differences in the relative proportions of certain annotated families. For example, *MAG_Ty3* content was proportionally greater than other lineages in MoO and *Copia_elem* content was proportionally greater than other lineages in MoS (Figure 1B). These differences in TE family prevalence across lineages hinted at complex lineage-specific TE dynamics, rather than genome-wide contraction or expansion of all TE content.

**Figure 1:**
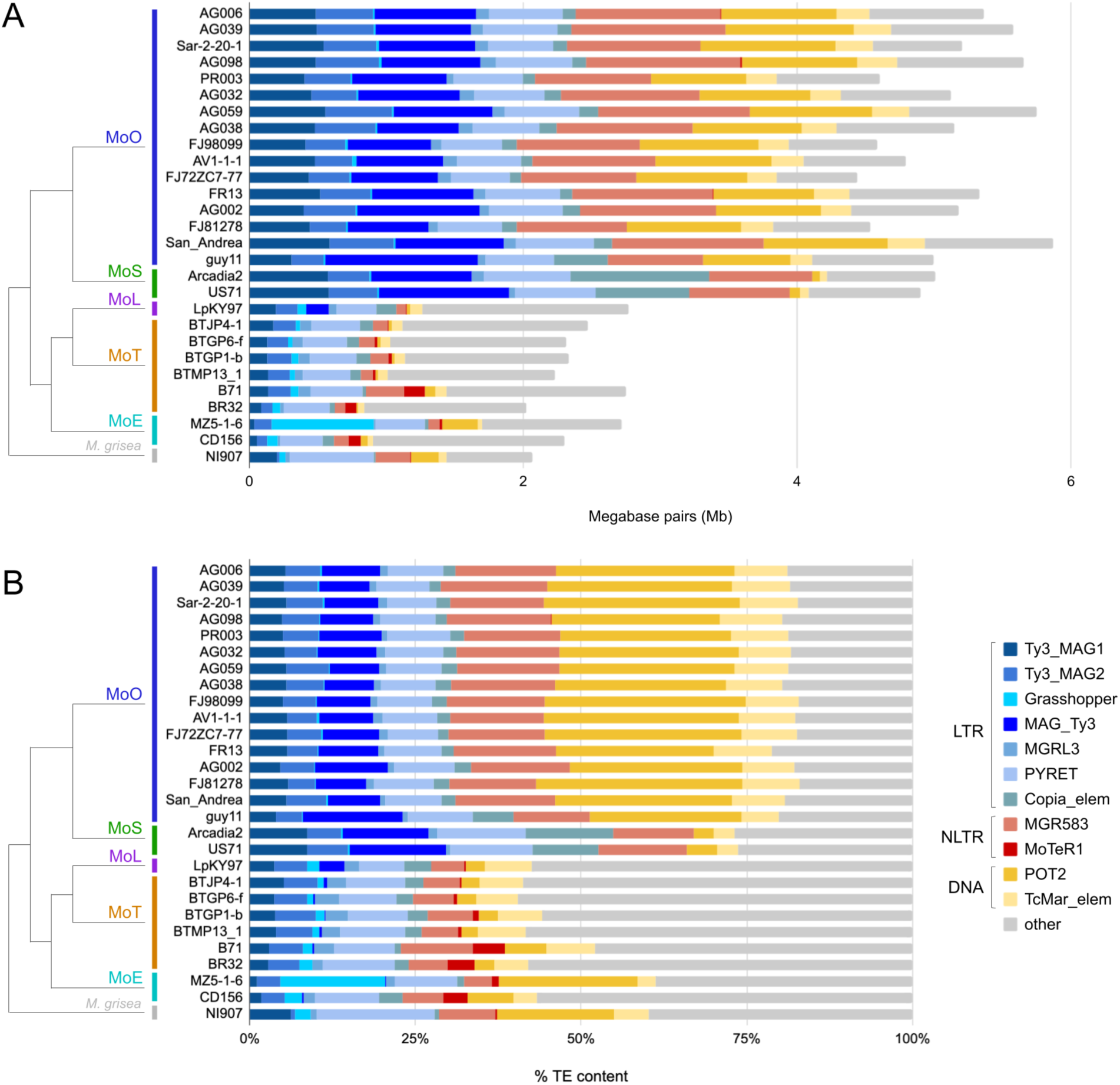
Large variation of TE content exists between *M. oryzae* genomes of different lineages. **A,** Stacked bar plot showing the number of megabase pairs (Mb) each TE occupies in each genome. **B,** Stacked bar plot showing the percentage that each TE family makes up out of all TEs in each genome. At the left of both plots is the lineage each genome belongs to, and the evolutionary relationships between lineages (Gladieux, Condon, et al. 2018) (branch lengths not to scale). Names of the TE families and their classification are shown in the key. LTR = long terminal repeat retrotransposon, NLTR = non-LTR retrotransposon, DNA = DNA transposon.

While there was an overall greater number of TEs in MoO and MoS, some families were more prevalent in the other lineages or in individual genomes. The *Grasshopper* LTR-retrotransposon made up a large portion of TE content in the MoE MZ5-1-6 genome specifically, but much less in the other MoE, MoT, and MoL genomes, and was absent in MoO and MoS. Additionally, the *MoTeR1* family had greater copy number in MoT’s B71 and BR32, and MoE’s CD156, but less in the other MoT, MoL, and MoE genomes, and was also absent from MoO and MoS (Figure 1B). This indicated that although MoT, MoL, and MoE have lower TE content, they may be more prone to isolate specific TE dynamics, in contrast to the larger and more uniform TE content in MoO and MoS.

### Recent lineage-specific expansions of LTR-retrotransposons led to differences in TE content between *M. oryzae* lineages

We next tested whether genome-wide or TE specific contraction or expansion dynamics led to the differences in TE content across lineages. Given that the bulk of the differences between MoO-MoS and other lineages seemed to be explained by LTR-retrotransposons (Figure 1B), we focused on these elements. We constructed domain-based maximum-likelihood (ML) phylogenies (Figure 2B-E, Figure S4) for each of the seven LTR-retrotransposons using all copies annotated in the highest quality genome of each lineage. The TE trees were compared to the genome tree (Figure 2A), which was generated based on the alignment of single copy orthologous genes (SCOs), in order to compare evolutionary relationships. Based on our analysis, three LTR-retrotransposon families stood out as having experienced lineage specific expansions. *MAG_Ty3* showed a large expansion in MoO and a smaller expansion in MoS (Figure 2B). *Copia_elem* expanded in both MoO and MoS (Figure 2C), and *Grasshopper* expanded only in the MoE MZ5-1-6 genome (Figure 2D). A helpful point of comparison was the *MGRL3* LTR-retrotransposon, which was present at a low copy number in all of the lineages. Elements in the *MGRL3* phylogeny didn’t strictly group by lineage and were more interleaved (Figure 2E), suggesting that it experienced an older expansion before the *M. oryzae* lineages diverged, and has not proliferated recently. The other LTR-retrotransposon phylogenies of *Ty3_MAG1*, *Ty3_MAG2*, and *PYRET* (Figure S4) also indicated older activity. These different histories show that a genome-wide deregulation of TEs was likely not responsible for the higher TE content in MoO and MoS, but rather that it resulted from TE family-specific dynamics.

**Figure 2:**
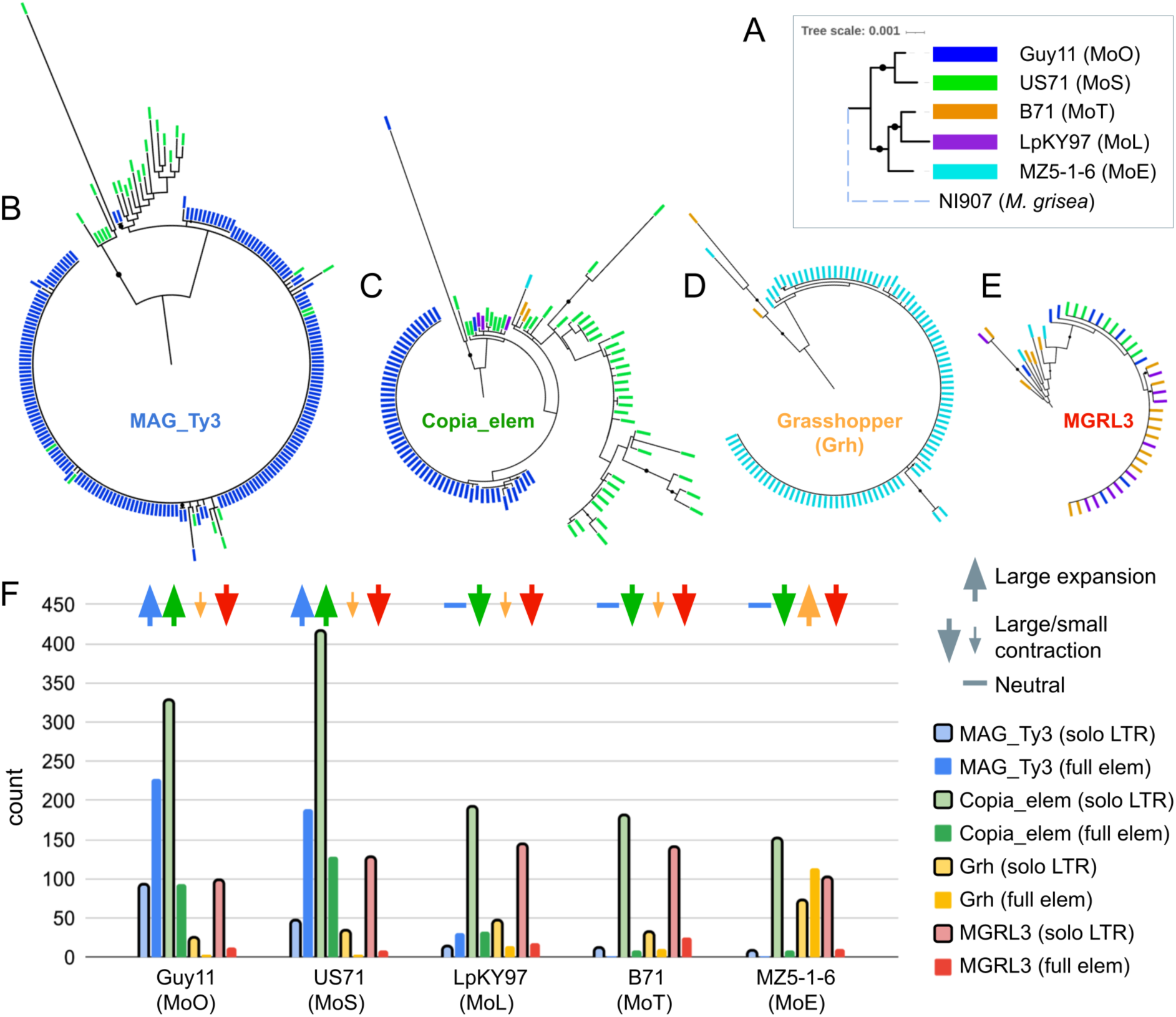
Certain LTR-retrotransposons have experienced lineage-specific expansions. **A,** Maximum-likelihood (ML) phylogeny of representative genomes of each lineage, based on the alignment of 8,655 SCOs. Branch lengths are to scale, except for the dashed line of the *M. grisea* outgroup. Bootstrap value of 100 is indicated by black circles. Domain-based ML phylogenies of TEs **B,** *MAG_Ty3*, **C,** *Copia_elem*, **D,** *Grasshopper*, and **E,** *MGRL3* are shown. Colored rectangle tips correspond to the genome each element is from, as shown in A, and black circles indicate bootstrap value ≥80. **F,** In this barchart, solo-LTR copy number is compared to the number of full length TEs to represent the expansion and contraction dynamics of TE families in each genome. Each bar is colored according to the specific family, corresponding to the color of the label within each TE phylogeny (B-E). Lighter bars outlined in black represent solo-LTRs. Arrows above each set of bars indicate our interpretation of the predominant explanation (expansion versus contraction) for the lineage-specific differences observed, based on the number of full elements versus solo-LTRs within the genome, and comparison of the number of full elements and solo-LTRs across all genomes.

We then wanted to see whether *Grasshopper*, *MAG_Ty3*, and *Copia_elem* had experienced lineage-specific expansions that occurred after all *M. oryzae* lineages had diverged, or whether expansions occurred in all lineages and were followed by subsequent losses. Since LTR-retrotransposons consist of an internal region that is flanked by direct repeats known as LTRs, they can be excised from the genome by non-allelic homologous recombination between the flanking LTRs (Wells and Feschotte 2020). When this occurs, a single LTR sequence known as a “solo-LTR” is left behind in the genome. We utilized the presence of these sequences to investigate TE expansion versus contraction dynamics. A large number of solo-LTRs and few full elements would suggest contraction of an LTR-retrotransposon population, while few solo-LTRs and many full elements suggest a recent expansion (Jedlicka et al. 2020). We observed that lineage-specific expansions are largely responsible for LTR-retrotransposon copy number variation, rather than the removal of these elements from some lineages (Figure 2F). *MAG_Ty3* had less than 13 solo-LTRs present in each of the MoL, MoT, and MoE lineages, which was much fewer than the 227 and 188 full-length *MAG_Ty3* in MoO and MoS, respectively. Thus, *MAG_Ty3’s* higher copy number in MoO and MoS was likely due to expansion in those lineages only. The *Copia_elem* family had a lot more solo-LTRs present in all genomes (>150), suggesting that older expansions may have occurred before the divergence of the lineages and that they were then partially removed. However, there were many more *Copia_elem* solo-LTRs in MoO and MoS (>300) along with more full copies, which could only have been achieved by expansions unique to MoO and MoS. *Grasshopper* had less than 50 solo-LTRs in all genomes besides MoE MZ5-1-6, which had 114 full-length elements, indicating an expansion in that genome only. In contrast, *MGRL3* had a relatively high number of solo-LTRs (>96) and a low number of full elements (<26) in all lineages. This supports the idea that *MGRL3* was expanded before the divergence of the lineages, then was largely removed from all of them over time. Thus, although both expansion and contraction play roles in determining LTR-retrotransposon copy number in *M. oryzae*, large expansions were the predominant cause of lineage-specific and isolate-specific copy number variation for the LTR-retrotransposons of interest.

### Complex LTR-retrotransposon proliferation history and dynamics explain lineage-specific expansions in *M. oryzae*

Next, we sought to better understand the timing and history of the LTR-retrotransposon expansions we observed. To do so, we used nucleotide sequence comparison and sequence divergence tests, which make the assumption that sequence divergence occurs at the same rate in all TEs. Thus, this assumption could be violated by the presence of repeat induced point mutation (RIP), a mutagenic mechanism in fungi that targets repetitive elements like TEs, causing GC to TA mutations (Pereira et al. 2021). RIP is only active during sexual reproduction (Ikeda et al. 2002), and previous studies have reported that it is minimally active in *M. oryzae* given its largely clonal life cycle (Ikeda et al. 2002; Pereira et al. 2021; van Wyk et al. 2021). However, there is evidence that RIP occurred during the large admixture event that potentially formed MoT and MoL (Rahnama et al. 2021), and RIP-associated genes *RID* and *DIM2* have been identified as present in some *M. oryzae* isolates (Bewick et al. 2019; van Wyk et al. 2021). An initial search of our orthogroups revealed that *RID* and *DIM2* were SCOs in our genomes, indicating the possibility of RIP affecting our TE divergence analyses. To measure how prevalent RIP was in our sequences, we calculated the GC content of all TE copies in each representative genome and compared them to the genome-wide average GC content of coding and non-coding regions (Pereira et al. 2021). We found that, although each TE family had a different median GC content, the individual copies did not deviate much from that value in most families (Figure 3). While we observed some trailing copies with low GC content, these were likely older elements that had been affected by RIP in the past. The *Pyret* LTR-retrotransposon had a GC content distribution with a strong skew that suggested it may have experienced RIP (Figure 3F). However, our data indicated that *Pyret* had not been recently active, since it had a similar number of copies in each lineage (Figure 1A) and elements in its phylogeny didn’t group by lineage (Figure S4), similar to *MGRL3*. Thus, RIP affecting Pyret had likely not occurred recently, and this family served as a good contrast to the recent LTR-retrotransposon expansions we focused on. As a final test, we compared the GC content of TEs in an MoO isolate originating from a recombining lineage (Guy11) to a clonal MoO isolate (FJ98099) (Latorre et al. 2020). There were no differences in the GC content between these two isolates, indicating very little RIP activity even in the sexually recombining MoO isolate (Figure S5). These analyses strongly suggest that although RIP may have been active in the past, it has had little effect on recently expanded TEs, and so the divergence tests we performed on our recently-expanded TEs were valid.

**Figure 3:**
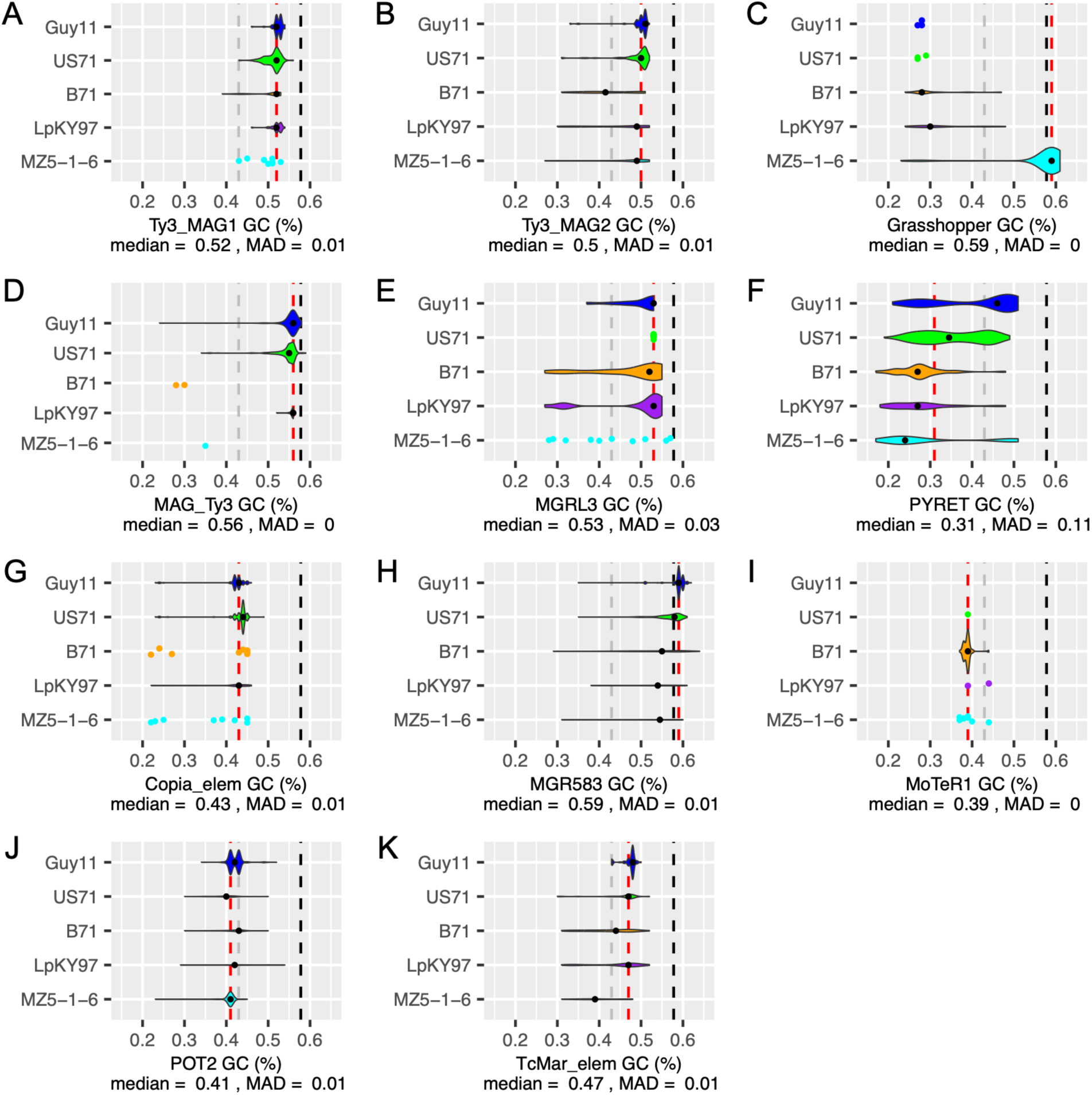
Repeat induced point mutation (RIP) has had little effect on recently expanded TE sequences. The GC content of each TE family in each representative genome is shown for **A,** *Ty3_MAG1*, **B,** *Ty3_MAG2*, **C,** *Grasshopper*, **D,** *MAG_Ty3*, **E,** *MGRL3*, **F,** *PYRET*, **G,** *Copia_elem*, **H,** *MGR583*, **I,** *MoTeR1*, **J,** *POT2*, and **K,** *TcMar_elem*. Data is shown as a violin plot unless there are <10 points, in which case a jitterplot was used. Within each plot, violin width is proportional to the number of TE copies represented. Dashed lines indicate: black = genome-wide average GC content of coding sequences, grey = non-coding sequences, red = the median GC content of the TE. The median GC content of the TE and the median absolute deviation (MAD) are specified.

To investigate the timing of LTR-retrotransposon expansions, we first sought to date individual TE insertions. A method for determining the age of LTR-retrotransposon insertions is to calculate the divergence between the flanking LTR sequences within an element (Jedlicka et al. 2020), since they are identical upon insertion (Wells and Feschotte 2020). Flanking LTRs of older elements would be more divergent, since they have had more time to accumulate mutations, while newer elements would have highly similar LTRs (Jedlicka et al. 2020). We determined LTR sequence divergence for *MAG_Ty3*, *Copia*, *Grasshopper*, and *MGRL3* retrotransposons (Figure 4A-D). Our results indicated that the expanded LTRs (*MAG_Ty3*, *Copia*, and *Grasshopper*) were inserted very recently, as many LTR pairs had zero sequence differences between them. The fact that LTR sequences are quite short (250 to 500 bp) combined with a reported mutation rate of 1.98e-8 substitutions/site/year in *M. oryzae* (Gladieux, Ravel, et al. 2018) likely contributed to this result. Nevertheless, given this mutation rate, a 500bp sequence would be expected to have mutated once in the past 50,000 years, indicating that these expansions could have occurred at any point since then, including more recently than the divergence of the MoO and MoS lineages 9,800 years ago (Gladieux, Ravel, et al. 2018). The *Copia_elem* in the MoS genome, on the other hand, showed a broader range of LTR divergence values, indicating proliferation events spread out over time (Figure 4B). *MGRL3* had slightly higher divergences between its flanking LTRs that were generally similar for all the lineages (Figure 4D), supporting the idea that it experienced an older expansion in a single period before the divergence of the lineages. Overall, these findings support our interpretation that the lineage specific LTR-retrotransposon expansions occurred recently.

**Figure 4:**
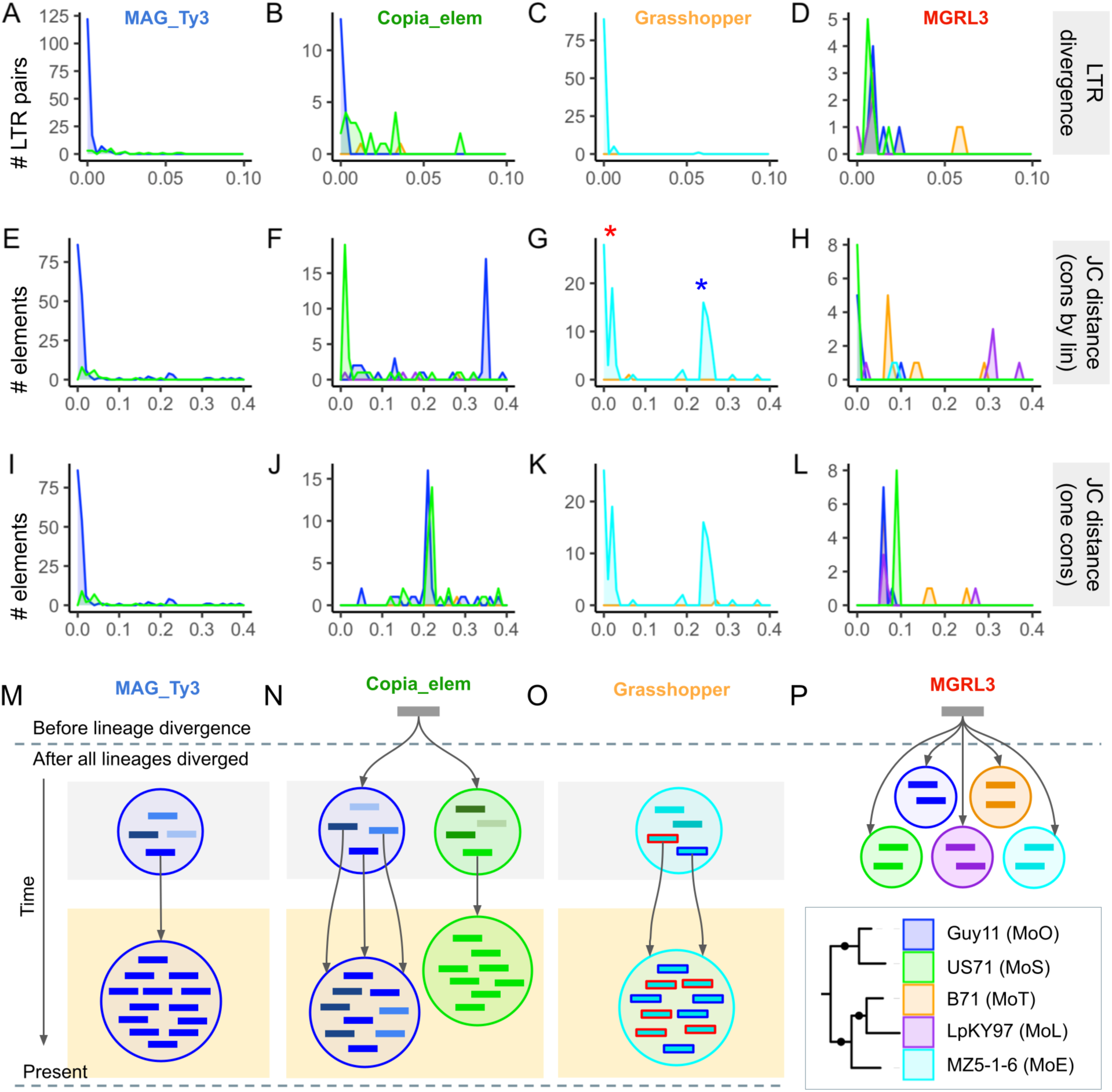
TE expansions in *M. oryzae* experienced complex histories that differ between various lineages. Columns correspond to *MAG_Ty3*, *Copia_elem*, *Grasshopper*, and *MGRL3* (from left to right, for each row of the figure). **A-D,** Divergence between flanking LTR sequences of LTR-retrotransposons. **E-H,** The Jukes-Cantor distance calculated using a separate consensus for each lineage. **I-L,** The Jukes-Cantor distance calculated using one consensus for all lineages. **M-P,** Schematic diagrams representing our hypothesis for the history of TE expansion events for each of the families. The representation of the current population for each TE is highlighted in yellow, and these expansions occurred either by one or multiple copies from an older population of TEs with sequence differences (highlighted in gray) proliferating recently. We indicate that TEs from different lineages might have proliferated from the same original copy for *Copia_elem* and *MGRL3*. Blue and red outlined *Grasshopper* rectangles correspond to the two labeled peaks in G. *MGRL3* is an example of an old expansion. The tree in the bottom right corner serves as a key for color-coding of the lineages for all parts of the figure.

Next, using the Jukes-Cantor distance metric, we estimated the sequence divergence of full-length TEs, following a previously published method (Faino et al. 2016). This analysis provided additional information beyond the TE phylogenies (Figure 2A-D), which were based only on the reverse transcriptase domain that each LTR-retrotransposon contains. For each TE, a consensus sequence was generated by aligning all copies of the family across all lineages. Then, the divergence of each copy from the consensus was determined and corrected by the Jukes-Cantor formula (Jukes and Cantor 1969). The same procedure was then repeated with a separate consensus for each lineage. The first method (one consensus for all lineages) showed the distance of each TE from the supposed common ancestor of that TE in all lineages, and indicated whether elements in each genome might have proliferated from the same original copy (Figure 4E-H). The second method (separate consensus for each lineage) showed how diverged the copies within one lineage were, and provided information on the recency of each expansion, and the population structure of TEs that contributed to it (Figure 4I-L). For *Grasshopper*, we found that the Jukes-Cantor distance metrics could have suggested two expansions of this family, one older and one more recent (Figure 4G). However, when taking into account our LTR divergence analysis (Figure 4C), it was more likely that the entire *Grasshopper* expansion occurred recently and consisted of multiple copies expanding in parallel (Figure 4O). We also looked at where the TE proliferations were localized in the genome and found that the *Grasshopper* expansion occurred globally regardless of the location of the original proliferating copy, as elements from both Jukes-Cantor peaks were distributed throughout MZ5-1-6’s seven chromosomes (Figure S6A). In contrast to *Grasshopper*, *MAG_Ty3* appeared to have expanded from a single element only (Figure 4E), but also very recently (Figure 4A). Our analyses of the *Copia* LTR-retrotransposon revealed a more complex scenario. Firstly, the family of *Copia* elements in MoO and MoS appeared to have proliferated from the same original copy, since most were about the same distance from the consensus of both lineages (Figure 4J). MoS *Copia_elem* copies were more similar to each other than those within MoO (Figure 4F). Yet, most MoO *Copia* had zero LTR divergence while many MoS *Copia* had further diverged LTRs (Figure 4B). The most likely explanation is that the *Copia* expansion in MoO occurred very recently, but consisted of multiple elements from an older *Copia* population with sequence differences. Meanwhile, the expansion in MoS was older and caused by just one copy proliferating (Figure 4N). Finally, *MGRL3* looked to have proliferated from the same original copy in all lineages (Figure 4L), which was consistent with the data supporting an old expansion of this family before the lineages diverged (Figure 2D,E). Overall, we have demonstrated that LTR-retrotransposons in *M. oryzae* have experienced complex proliferation dynamics, resulting in different histories of each lineage-specific expansion.

### A DNA transposon, *POT2*, appears to have been transferred from the rice pathotype of *M. oryzae* to the wheat pathotype

Although LTR-retrotransposons had a greater role in increasing the TE content of MoO and MoS, the *POT2* DNA transposon also stood out as being a large contributor (Figure 1). As shown by the ML phylogeny based on alignment of *POT2’s* transposase domain, we found it to have greatly expanded in MoO and MoE, with a smaller expansion in MoT (Figure 5A). The phylogeny also suggested a potential transfer of *POT2* between MoO and MoT due to the unexpectedly high similarity of certain copies from MoT B71 and MoO Guy11. Previously published criteria for identifying potential TE horizontal transfers (HTs) based on phylogenies are: (i) unexpectedly high similarity between TEs in lineages that aren’t closely related, (ii) a patchy distribution of the element in one of those lineages, as well as absence from its sister lineage, and (iii) discordance between the TE tree and genome tree (Bergman 2018). When comparing the *POT2* phylogeny to the genome tree based on SCOs (Figure 2A), we observed clear discordance between the two. We expected MoT *POT2* to be more closely related to MoE *POT2*, since MoT is closer to MoE than to MoO but this was not the case. Additionally, *POT2* copies from another MoT genome (BR32) were not found in the clade containing potentially transferred *POT2* from B71 (Figure S7). BR32 *POT2* copies were also not found to be expanded, likely indicating a patchy distribution of *POT2* in MoT. Finally, *POT2* was generally absent from MoL, the most closely related lineage to MoT. Although there were some older copies of *POT2* in all genomes, including MoL, no *POT2* from MoL were found in or near the clade containing potentially transferred MoT *POT2* (Figure 5A). These results are in line with the criteria for a HT of *POT2*.

**Figure 5:**
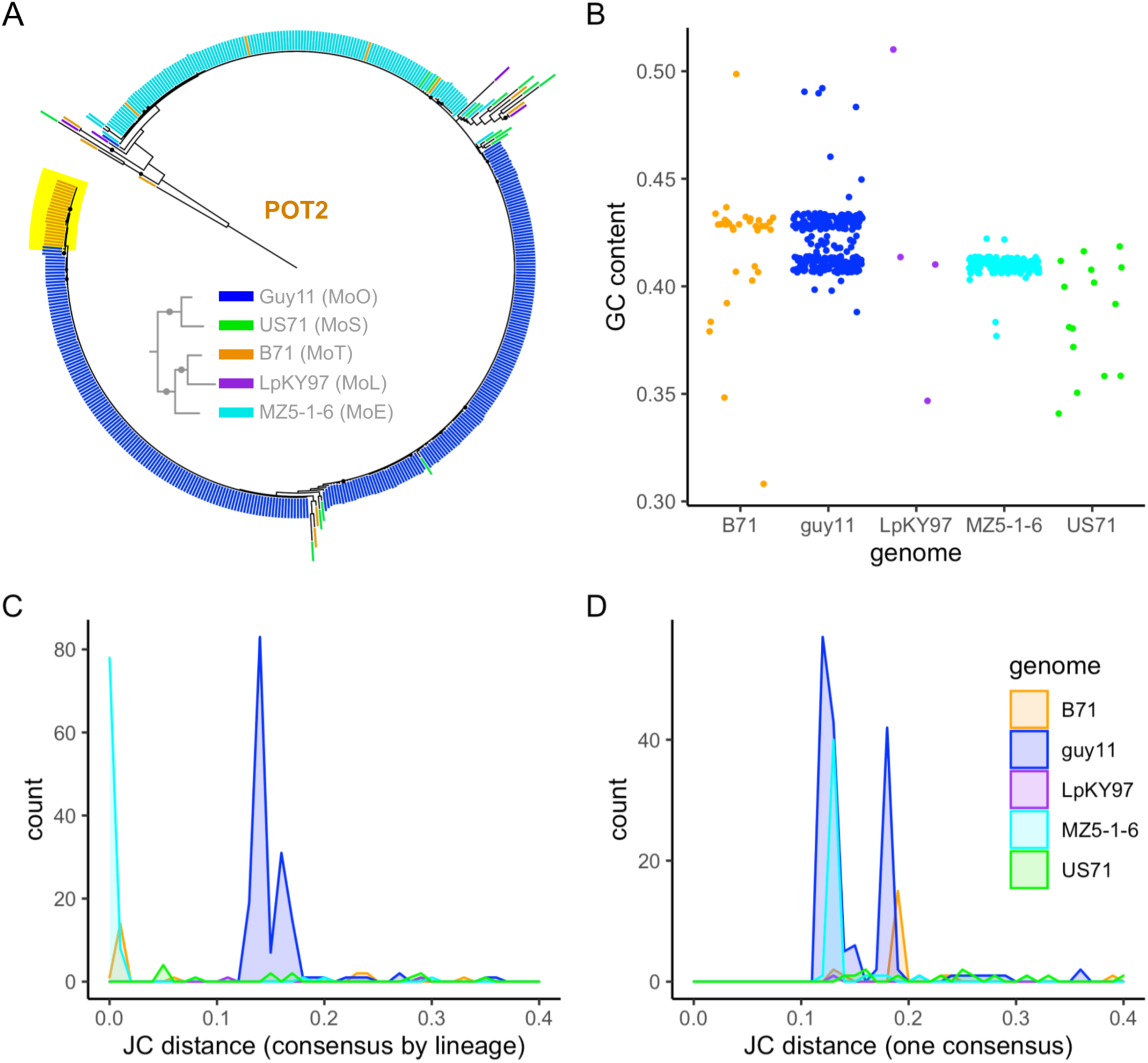
*POT2* was likely transferred between MoO and MoT. **A,** Domain-based ML phylogeny of *POT2* in each representative genome (Guy11, US71, B71, LpKY97, MZ5-1-6). The yellow highlighted MoT elements indicate those that were potentially transferred, and black circles indicate bootstrap value ≥80. The smaller phylogeny within shows the expected relationships between the lineages for comparison (same as Figure 2A) and the color coding, representing the lineage an element is from. **B,** Jitter-plot showing GC content in each *POT2* copy, in each genome. **C,** Jukes-Cantor distance analysis of *POT2* based on a separate consensus for each lineage. **D,** Jukes-Cantor distance analysis of *POT2* based on one consensus for all lineages.

We also observed additional lines of evidence pointing to a potential transfer of *POT2* between MoO and MoT. Our analysis of GC content revealed that the MoO Guy11 genome had two distinct groupings of *POT2* resembling subfamilies, one with higher GC content and one with slightly lower GC content (Figure 5B). Many *POT2* in MoT had the same higher GC content as the former subfamily, and most *POT2* in MoE had the same lower GC content as the latter. The difference in GC content between the *POT2* subfamilies could have been caused by the ancestor of the MoO-MoE *POT2* being slightly RIPped, while the MoO-MoT *POT2* ancestor had not, then each element having had its own evolutionary trajectory thereafter. The Jukes-Cantor analysis further supported the trend observed from the GC content analysis. When comparing *POT2* copies from all lineages to their consensus, we saw the same two MoO-MoT and MoO-MoE subfamily groupings (Figure 5D). This supported the idea that the two groups didn’t come from the same original *POT2* element. Instead, they likely originated from separate elements with sequence differences. Although the MoO-MoE grouping of *POT2* by GC content and Jukes-Cantor distance might resemble a transfer between these lineages as well, it is not supported by the phylogeny (Figure 5A). Thus, the original MoO-MoE *POT2* was likely present in all lineages but only expanded in MoO and MoE, while the MoO-MoT *POT2* expanded only in MoO then transferred to MoT. Comparing each *POT2* copy to the consensus of its lineage (Figure 5C) showed that *POT2* in MoT and MoE were more closely related within their respective lineages than *POT2* within MoO. This suggested that either *POT2* expansions in MoT and MoE occurred much more recently than in MoO, or that they consisted of a single element expanding, while multiple elements from a population of TEs with sequence differences expanded in MoO. Either interpretation supported the idea that an individual *POT2* was recently transferred from MoO to MoT and subsequently expanded.

*POT2* also experienced differential localization of its expansions in different lineages. Local proliferation was displayed by *POT2* in MoT, where most of its copies were located on the minichromosome sequences of the B71 genome (Figure S6B). In contrast, *POT2* in MoE was evenly distributed throughout the seven chromosomes (Figure S6C), which was similar to the LTR-retrotransposon expansions we characterized. Minichromosomes have been reported to harbor many repetitive sequences as well as virulence factors (Langner et al. 2021). Of the genomes used in this study, it is known that MZ5-1-6 (MoE), BR32 (MoT), and Guy11 (MoO) do not have minichromosomes, while B71 (MoT), LpKY97 (MoL), FR13 (MoO), US71 (MoS), and CD156 (MoE) do (Peng et al. 2019; Langner et al. 2021). Despite the existence of genomes both with and without minichromosomes in many of the lineages, their presence did not affect the lineage-specific patterns of TE content (Figure 1). Since the MoE MZ5-1-6 genome does not contain minichromosomes and experienced a global *POT2* expansion, while B71 does have minichromosomes and had a local *POT2* expansion there, it is possible that the presence of minichromosomes affects the overall localization of TE content in *M. oryzae*. Regardless, this result further highlights the different expansion histories and dynamics of *POT2* in different lineages.

### *POT2* may have been transferred during a recombination event between isolates of the rice and wheat *M. oryzae* pathotypes

To investigate how *POT2* may have been transferred between the MoO and MoT lineage, we first looked for any regions of the Guy11 and B71 genomes that may have been involved in a larger transfer event. *POT2* elements and their flanking regions were compared using DNA alignments and synteny analysis; however, none of these segments containing *POT2* stood out as being potentially transferred regions. We then considered the possibility that *POT2* could have moved as part of a larger region but then transposed out of that region. We looked for evidence of genes that might have been transferred between Guy11 and B71 by filtering for gene trees that followed the same topology as the *POT2* phylogeny. One region in B71 on chromosome 7 stood out as having many of these genes, with 29 out of the 38 genes that matched the *POT2* phylogeny being located there (Figure S8A). The other 9 genes were scattered among various chromosomes. We located the orthologs of the 29 B71 genes in the other genomes, and found them to be syntenic. In LpKY97 and MZ5-1-6, the other two chromosome level assemblies, the genes were in the same location on chromosome 7. We aligned the full-length nucleotide sequence from each genome and produced an ML phylogeny (Figure 6A), which showed that this entire region followed a *POT2*-like tree topology rather than the expected evolutionary relationships between the *M. oryzae* lineages (Figure 2A). Since B71 grouped with Guy11 in the MoO-MoS clade, it was likely that an MoO isolate was the donor of this region in B71. Additionally, since this region was syntenic in all genomes, the most likely explanation for the transfer event was homologous recombination.

**Figure 6:**
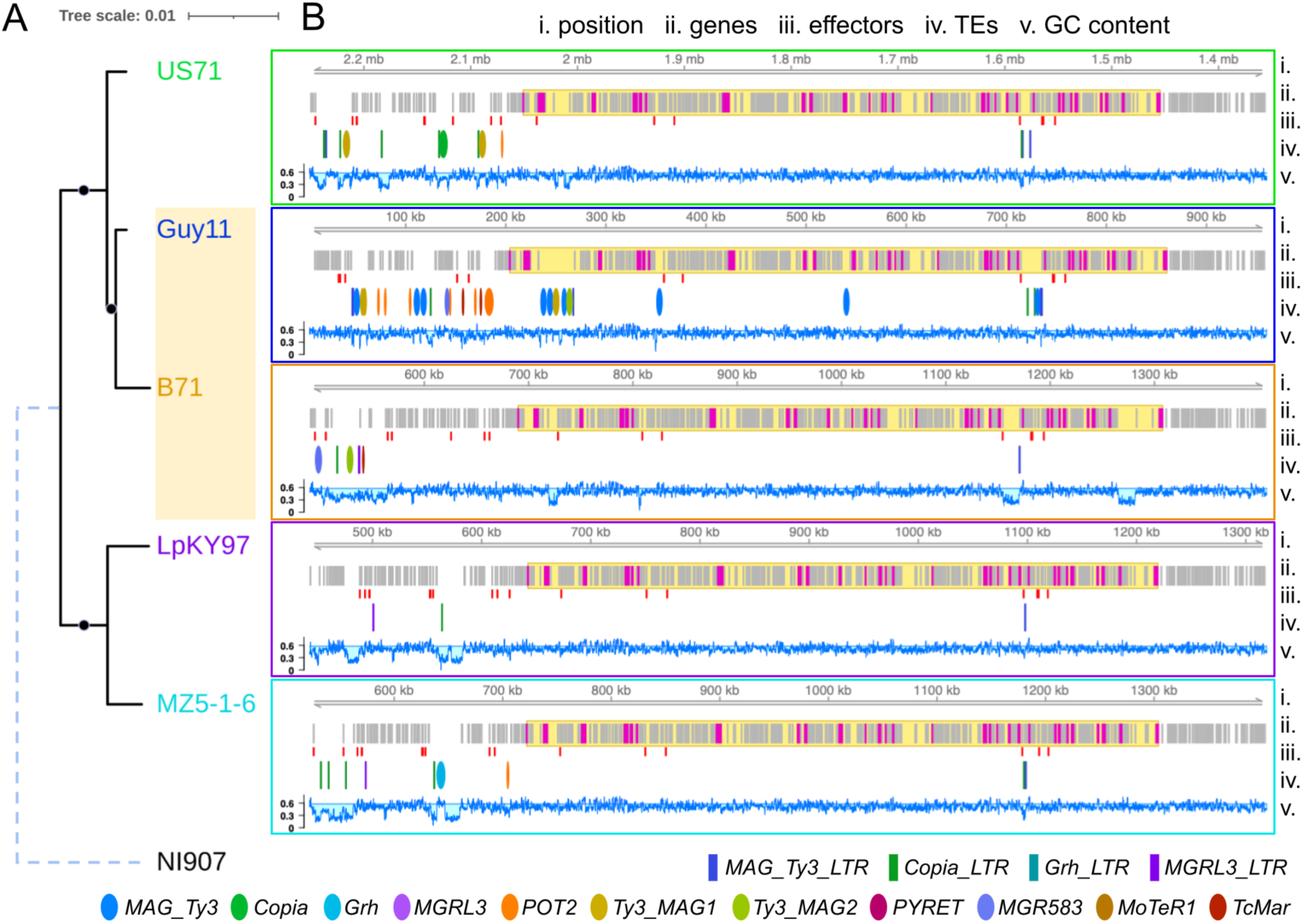
A large region may have been transferred between MoO and MoT isolates as a result of recombination. **A,** The ML phylogeny of an ∼583 kb syntenic region on chromosome 7 that contains many genes following a *POT2*-like gene tree topology. Black circles indicate a bootstrap value of 100. Branch lengths are to scale, except for the dashed outgroup branch of NI907 (*M. grisea*). **B,** Genomic tracks show features of the potential region of recombination in each genome. Tracks from top to bottom: i) Position along the scaffold or chromosome (B71: CM015706.1, Guy11: MQOP01000008.1, US71: UCNY03000007.1, LpKY97: CP050926.1, MZ5-1-6: CP034210.1, NI907: CM015044.1). ii) All genes, where magenta represents a *POT2*-like topology gene, and the yellow highlighted area indicates the region containing all of those genes. iii) Position of candidate effectors. iv) Position of TEs, where ellipses are full elements (blue=*MAG_Ty3*, green=*Copia_elem*, teal=*Grasshopper*, purple=*MGRL3*, orange=*POT2*, mustard=*Ty3_MAG1*, yellow-green=*Ty3_MAG2*, magenta=*PYRET*, skyblue=*MGR583*, light-brown=*MoTeR1*, tomato=*TcMar_elem*) and rectangles are solo-LTRs (dark-blue=*MAG_Ty3_LTR*, dark-green=*Copia_LTR*, dark-teal=*Grasshopper_LTR*, dark-purple=*MGRL3_LTR*). v) GC content, where the horizontal line is the genome-wide average GC content of coding regions (0.577891).

We then looked at the TE insertions in the region we identified to determine the timing of the TE expansions we characterized in relation to the transfer of the region. There were no full length TEs contained in this region besides in the Guy11 genome, where *MAG_Ty3*, *Ty3_MAG1*, and *Ty3_MAG2* elements were likely inserted after the transfer event. There were a few solo-LTRs located in the region, including a *MAG_Ty3* solo-LTR that was present at the same location in all genomes. Located upstream of the transferred region there was a unique set of many TEs in each genome, indicating lineage-specific TE activity. There were no *POT2* within or nearby this region in B71, however there were many *POT2* copies upstream of the region in Guy11 (Figure 6B), supporting the possibility that one of these elements were included in the recombination event.

Finally, we investigated if any genes of importance were transferred along with this region. There were a few predicted effectors (Figure 6B), however they were not under presence-absence variation and did not include any AVRs or members of expanded *M. oryzae* ART and MAX effector families (Seong and Krasileva 2021). We then characterized the genes in this region by obtaining their Gene Ontology (GO) terms (Additional File 2) and PFAM domain terms (Additional File 3). The most common terms included a putative ssRNA binding PFAM domain (RRM_1), iron ion binding molecular function (MF), zinc ion binding MF, proteolysis biological process (BP), glycolytic process BP, DNA repair BP, and mitochondrion cellular component (CC). Although the region was too small (182 genes out of 12,658 total) to perform meaningful enrichment analysis, it notably had similar characteristics to a recently discovered group of large mobile elements known as *Starships*, due to the presence of effectors, metal-binding genes, and other conserved genes (Gluck-Thaler et al. 2022). Within the region, there were genes containing NOD-like receptor (NLR)-associated domains (HET, Ank_2, Ank_4, Ank_5) and ferric reductase-associated domains (FAD_binding_4, FAD_binding_7, NAD_binding_2, NAD_binding_11), both of which are conserved genes in *Starships* (Gluck-Thaler et al. 2022). While tyrosine recombinase (DUF3435) and patatin-like phosphatase *Starship*-associated domains were not present within the region, a nearby upstream gene in B71 contained a fragmented DUF3435 (Figure S8B). It is possible that this region could have originated from a *Starship* element, however it would likely be much older than lineage divergence, given the synteny in all genomes. Thus, our results still strongly suggest that recombination caused the transfer of the region. Yet, it is interesting to consider the potential origins of the region, due to the adaptive function often conferred by mobile *Starship* elements (Gluck-Thaler et al. 2022).

## Discussion

Differences in TE content contribute to intraspecific diversity in many fungal species (Castanera et al. 2016; Shirke et al. 2016; Chen et al. 2018; Bleykasten-Grosshans et al. 2021; Oggenfuss et al. 2021; Gourlie et al. 2022). To understand how TE content variation arises and may relate to the evolutionary history of fungal pathogens, we constructed a *de novo* TE library that represents the diversity of TEs in five *M. oryzae* lineages. Using this library, we found that MoO and MoS contain much greater TE content than MoT, MoL, and MoE. Our pipeline ensured that these differences were not due to any bias of the TE library towards particular lineages, and correlation tests showed that they were not due to differences in genome assembly or quality. While we focused on elements containing TE-associated domains (Muszewska et al. 2019), further work is needed to study the full set of all repetitive DNA across the *M. oryzae* lineages. Most notably, protein domain-lacking elements such as MITEs are not included here, so our analyses likely underestimate overall TE content. Additionally, our analysis was restricted to highly contiguous genome assemblies which are few in number for *M. oryzae*, especially for the MoS, MoL and MoE lineages. While we are confident that our analyses are robust for the TE families we characterized, we may have missed other isolate-specific TE expansion events.

Despite these limitations, we found strong evidence that recent lineage-specific TE expansions contributed to the greater number of TEs in MoO and MoS. Analyzing solo-LTR copy numbers allowed us to verify that some LTR-retrotransposons were expanded in certain lineages, rather than having been expanded in all lineages and subsequently removed in only some. By synthesizing the results of our LTR divergence and Jukes-Cantor distance analyses, we were able to construct a model showing differences in the method of TE expansion between various types of TEs and between the same TE in different lineages. Some expansions were caused by the proliferation of one element, while others consisted of multiple elements from an older population of TEs with sequence differences proliferating in parallel. Most expansions occurred globally, with elements being distributed throughout the genome, however the *POT2* DNA transposon proliferated locally in the MoT isolate (B71) minichromosomes. Solo-LTR and LTR divergence analyses are not possible for DNA transposons, so it is difficult to determine expansion versus contraction dynamics for *POT2*, and how recently its copies proliferated. Nevertheless, our reconstruction of TE expansion histories points to the complexity of TE activity in *M. oryzae*.

Through our analyses, we found multiple lines of evidence suggesting the recent transfer of a DNA transposon between rice and wheat-infecting *M. oryzae* lineages. The phylogeny, Jukes-Cantor distances, and GC contents of *POT2* copies all showed that MoT *POT2* grouped unexpectedly with MoO *POT2* when considering the evolutionary relationships between the lineages. Given that *POT2* has been found to insert into the *AVR-Pib* effector gene in MoO field isolates and modulate their virulence (Li et al. 2022), its transfer to other lineages has the potential to contribute to their adaptability. The potential transfer of *POT2* could have occurred by a variety of mechanisms, including HT or recombination between the lineages. Notably, *POT2* is a DDE-type DNA transposon of the Tc1/Mariner family, which are reported to be prone to HT (Wells and Feschotte 2020). However, we could not identify direct evidence of such an HT event. It is possible that an individual *POT2* transferred by itself, which would not be possible to detect through a comparative genomics approach, given that DNA transposons leave almost undetectable excision footprints (Luo et al. 1998). Additionally, we did not find evidence of any non-syntenic, horizontally transferred regions that could have carried *POT2.* Since the potentially transferred *POT2* copies are localized on B71’s minichromosomes, it is also possible that minichromosome dynamics allowed *POT2’s* transfer or resulted in its localization. The HT of minichromosomes between isolates has been previously observed (Langner et al. 2021), as has the acquisition of core chromosomal regions by minichromosomes (Peng et al. 2019). An alternative explanation for the transfer of *POT2* is gene flow between the lineages through recombination during sexual reproduction. A previous study has shown evidence of historical gene flow in *M. oryzae*, most of which was caused by events that occurred before the divergence of the lineages (Gladieux, Condon, et al. 2018). However, the hypothesis that MoT and MoL arose within the last 60 years via a large admixture event involving recombination between isolates of various pathotypes in South America (Rahnama et al. 2021) makes it possible that *POT2* was acquired by MoT in that event.

In searching for genes that might have accompanied *POT2* in a potential transfer event, we identified a region of recombination between MoO and MoT. This region on chromosome 7 contained many genes whose phylogenies followed the topology of the *POT2* phylogeny, and were syntenic in each lineage. Although our analyses don’t rule out the possibility of incomplete lineage sorting, this region was also identified by Rahnama *et al*. as originating from a currently unsampled (cryptic) relative of MoO and MoS (Rahnama et al. 2021) that participated in recombination during the large admixture event. This likely occurred before the divergence of MoT and MoL, since a few MoL isolates appear to contain the region and a few MoT isolates do not (Rahnama et al. 2021). This strongly suggests that the region we identified experienced recombination. The TE insertions in this region in the MoO isolate provide further evidence that the transferred region originates from a relative of MoO that participated in the admixture event, since it’s unlikely that this region accumulated eight new LTR-retrotransposon insertions in the past 60 years (Figure 6B). Since we did not find a copy of *POT2* in the transferred region, there was no direct evidence that POT2 transferred by this mechanism. However, the exact boundaries of the transferred region are unclear, so it’s possible that *POT2* copies present upstream in MoO Guy11 could have been included. Thus, we highlight this region as an example of a potential way that *POT2* may have been transferred. We propose that *POT2* was transferred from the cryptic relative of MoO and MoS to an ancestor of MoT and MoL in the recombination event involving this region. Subsequently, *POT2* could have transposed out of the transferred region to B71’s minichromosomes, where it proliferated (Figure 7).

**Figure 7:**
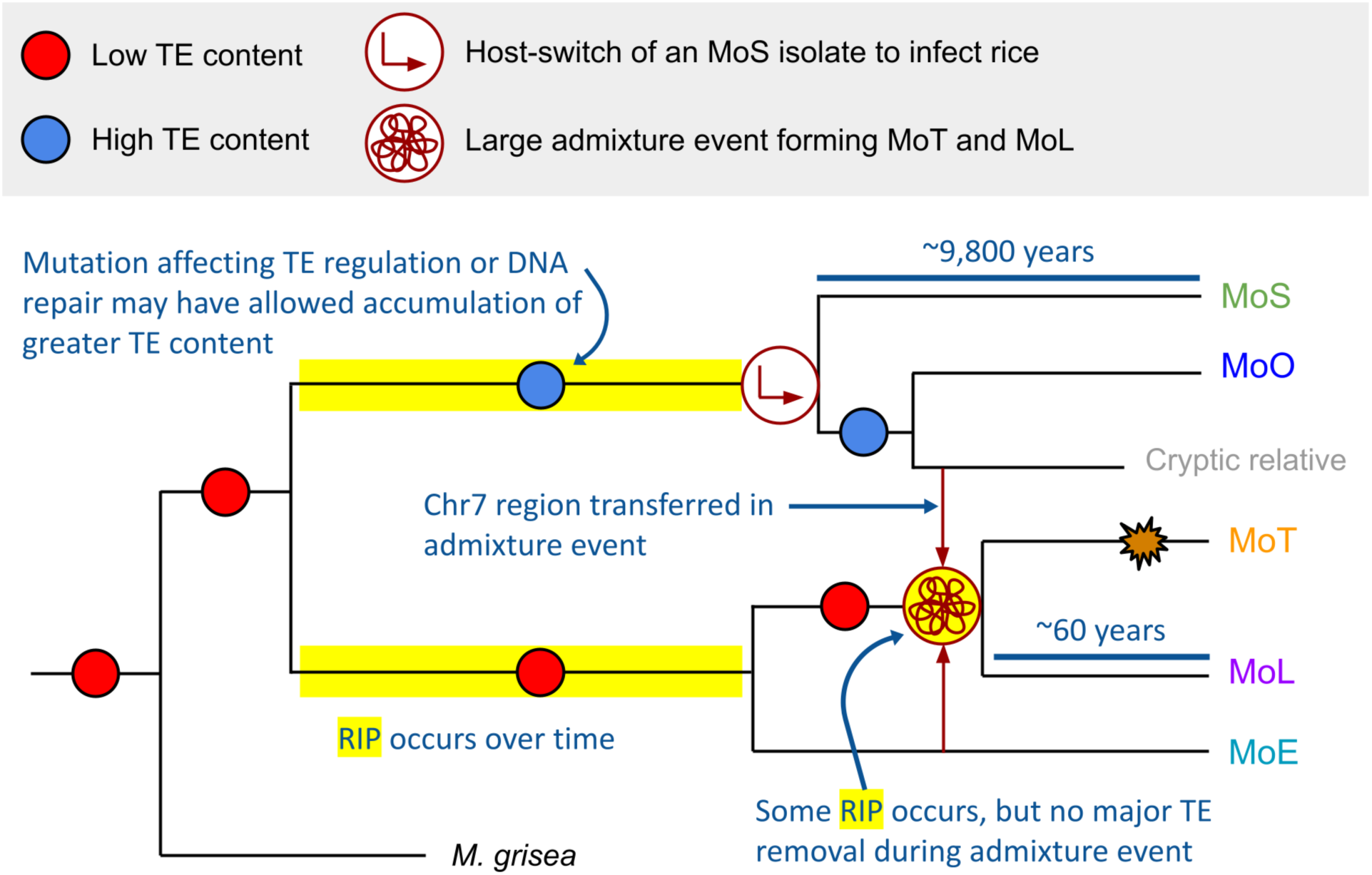
Proposed model showing how the MoO-MoS and MoT-MoL-MoE lineage groups have likely experienced different evolutionary histories. MoO-MoS divergence via host switch is well accepted, while our results indicate that MoT and MoL were formed in a large admixture event, rather than as a result of a conventional host switch. The diversity present in MoO-MoS accumulated over approximately 9,800 years, while the diversity in MoT-MoL was generated over the past 60 years. Our results suggest that a lower TE content was likely the ancestral state. We hypothesize that some difference, possibly a mutation affecting a TE regulation or DNA repair pathway, may have contributed to the increased TE content in MoO and MoS. RIP (highlighted in yellow) may have occurred during the large admixture event forming MoT and MoL, but is not sufficient to explain the higher TE content in MoO and MoS. During the admixture event, a region on chromosome 7 (Chr7) was transferred from a cryptic relative of the MoO lineage and may have contained a *POT2* element, which subsequently expanded in the MoT lineage (orange starburst).

Taken together with previous findings, our results suggest that the MoO-MoS and MoT-MoL-MoE lineage groups have experienced distinct evolutionary histories. MoO and MoS are set apart by their greater TE content; although the TE expansions we characterized help to explain some of the difference, it is still unclear how they accumulated approximately twice as many TEs as the other lineages. We hypothesize that low TE content was the ancestral state, and perhaps a mutation causing a difference in a TE regulation or DNA repair mechanism in the ancestor of MoO-MoS allowed the accumulation of greater TE content over time (Figure 7). Genes involved in DNA repair are of particular interest due to the recent finding that multiple non-canonical and error-prone DNA repair pathways exist in *M. oryzae*, and their influence on genomic variation are not well understood (Huang et al. 2022). Notably, TEs in *M. oryzae* have been found to activate in response to stress (Chadha and Sharma 2014), so various environmental and host-plant conditions may have also played a role in TE content differences. MoT, MoL, and MoE are distinguished by their greater isolate-specific TE content variation (Figure 1), and multiple lines of evidence suggest that MoT and MoL were the result of a large admixture event (Rahnama et al. 2021), rather than a host-switch of MoL to infect wheat as previously hypothesized (Inoue et al. 2017). This is in contrast to MoO and MoS, which are well established to have diverged via a host-switch of MoS to rice (Couch et al. 2005; Gladieux, Ravel, et al. 2018; Zhong et al. 2018). The two hypotheses represent quite different evolutionary processes, as a host-switch implies steady adaptation and selection for mutations that allow infection of a new host, while a large admixture event may quickly bypass the process of adaptation by generating lots of diversity, causing some strains to infect new hosts. Along with the argument presented by Rahnama *et al*. (Rahnama et al. 2021), our findings of the potentially transferred *POT2* and region of recombination align with the admixture hypothesis for MoT and MoL. Finally, although the diversity represented in the genome tree by branch lengths is comparable for the two lineage groups (Figure S2), it’s important to note that the diversity we observed was likely generated much more rapidly for MoT-MoL in the last 60 years (Rahnama et al. 2021), compared to MoO-MoS, where it seems to have accumulated steadily over 9,800 years (Gladieux, Ravel, et al. 2018). A large admixture event recently forming MoT and MoL, as compared to the host-switch resulting in the divergence of MoO and MoS, would explain these differences well.

In formulating our model (Figure 7), we considered whether the large admixture event presented by Rahnama *et al*. (Rahnama et al. 2021) might explain the low TE content of MoT and MoL, where many TEs could have been mutated beyond recognition by RIP and removed via recombination during the event. However, our data supports the alternative hypothesis that all lineages originally had lower TE content, and MoO-MoS independently accumulated their greater TE content after divergence from the common ancestor of all lineages (Figure 7). MoE genomes have low TE content, so this is not unique to the MoT-MoL isolates that originated from the admixture event. Likewise, the fact that TE content of the *M. grisea* outgroup is most similar to MoT-MoL-MoE supports the idea that the common ancestor of all lineages had low TE content. Additionally, if many TEs were removed during the admixture event, we’d expect large numbers of solo-LTRs in MoT and MoL, since recombination between flanking LTRs is a common mechanism of removal. However, our results indicate no extensive removal in MoT and MoL to explain the large difference in TE content. A lack of severe RIP in the TEs we analyzed also refutes the idea that RIP mutated TEs beyond recognition in the genomes of all lineages except MoO and MoS. Through our GC content analysis (Figure 3, Figure S5) we concluded that there had not been recent RIP affecting TEs in any *M. oryzae* lineages. It is very likely that RIP was active during the admixture event since there were many sexual recombinations, and Rahnama *et al*. found regions that had clearly been RIPped (Rahnama et al. 2021). However, this RIP activity was not sufficient to explain the differences in TE content we observed. The strongest evidence showing that RIP had not severely affected TEs during the admixture event is that the *MGRL3* LTR-retrotransposon, which we found to be an old element that proliferated before the divergence of the lineages, had not experienced recent RIP in any of the lineages (Figure 3). If RIP affected *MGRL3* during the admixture, we would expect to find less copies in MoT and MoL compared to other lineages. However, its copy number of full elements and solo-LTRs was very uniform throughout all lineages. Thus, we propose that there is a biological difference between the MoO-MoS and MoT-MoL-MoE clades responsible for the drastic difference in TE content, likely related to their different evolutionary histories (Figure 7).

In this study, we have shown that lineage-specific TE content differences in *M. oryzae* were caused in part by complex lineage-specific TE expansion dynamics. Future studies might investigate potential DNA repair or TE regulation mechanisms to better understand how the large differences between the MoO-MoS and MoT-MoL-MoE lineage groups arose. Additionally, the dynamics of a DNA transposon potentially transferred from MoO to MoT led us to identify a region of recombination between the two lineages. Our results support the hypothesis that the MoT and MoL lineages were formed in a large admixture event rather than caused by a host-switch, suggesting that their evolutionary history was vastly different from MoO and MoS. Further work is needed to investigate the consequences of differing evolutionary histories on the adaptive potential and trajectory of *M. oryzae* lineages. This study demonstrates that investigating TE dynamics can help us to better understand intraspecific diversity, which is especially important in fungal pathogens with host-specific populations.

## Methods

### Genomic datasets used and quality assessment

All genome sequences were retrieved from NCBI GenBank in December 2020, along with information on the host they were isolated from, the year they were collected, their GenBank accession, the assembly quality, the number of scaffolds, and the genome size (Table S1). Isolates were chosen primarily based on having the lowest number of contiguous scaffolds, which is ideal for TE annotation (Rech et al. 2022). We assessed the completeness of the genomes using BUSCO (Manni et al. 2021) version 5.2.2 software with ‘sordariomycetes_odb10’ as the busco_dataset option.

### TE annotation, classification, and phylogenetic analysis

The highest quality representative genomes of each lineage (Guy11 for MoO, US71 for MoS, B71 for MoT, LpKY97 for MoL, and MZ5-1-6 for MoE) were used as input into Inverted Repeat Finder (Warburton et al. 2004) version 3.07 and RepeatModeler (Flynn et al. 2020) version 2.0.2 software to obtain *de novo* annotations of TEs representing all lineages. Inverted Repeat Finder was called with options ‘2 3 5 80 10 20 500000 10000 -a3 -t4 1000 -t5 5000 -h -d -ngs’, and RepeatModeler was called with options ‘-engine ncbi -LTRStruct’. These libraries were combined with the RepBase (Bao et al. 2015) fngrep version 25.10 library of known TEs in fungi, and clustering was performed using cd-hit-est from CD-HIT (Fu et al. 2012) version 4.7 with options ‘-c 1.0 -aS 0.99 -g 1 -d 0 -M 0’. The resulting comprehensive TE library was then scanned against a list of domains containing TE coding sequences (Muszewska et al. 2019), consisting of both CDD profiles and PFAM profiles. The TE library was scanned for CDD profiles using rpsblastn from BLAST (NCBI Resource Coordinators et al. 2018) version 2.7.1+ with option ‘-evalue 0.001’. PFAM profiles were retrieved from the Pfam-A.hmm file included with HMMER (Eddy 2011) version 3.1b2. PFAM hmms were used for domain scanning along with pfam_scan.pl (Finn et al. 2016) version 1.6 with options ‘-e_dom 0.01 -e_seq 0.01 -translate all’. Elements not containing at least one of either a PFAM or CDD domain were filtered out.

Next, new sequences generated from *de novo* repeat-finding were classified in the library. Domain-based ML phylogenies were constructed using the most common PFAM domains found in the TE library, which were RVT_1 (PF00078.29), DDE_1 (PF03184.21), rve (PF00665.28), Chromo (PF00385.26), RNase_H (PF00075.26), and RVT_2 (PF07727.16). Each domain was aligned to the TE library using HMMER hmmalign (Eddy 2011) version 3.1b2 with options ‘--trim --amino --informat fasta’. The alignments were processed using esl-reformat and esl-alimanip from Easel version 0.48, which is part of the HMMER (Eddy 2011) package. Columns containing all gaps were removed by calling esl-reformat with the ‘--mingap’ option, so that the length of the alignment was the same as the hmm length. Then, sequences that didn’t match at least 70% of the hmm were filtered out by calling esl-alimanip with the ‘--lmin’ option specifying 70% of the hmm length. Finally, esl-reformat was used to convert the alignment to fasta format. RAxML (Stamatakis 2014) version 8.2.11 with options ‘-f a -x 12345 -p 12345 -# 100 -m PROTCATJTT’ was then used to construct the domain-based ML phylogeny of TEs containing the domain. This process was repeated for each of the six domains. The phylogenies were visualized in the Interactive Tree of Life (iTOL) (Letunic and Bork 2019) online tool, and clades where *de novo* elements grouped with elements having a classification in the RepBase library were copied using the “copy leaf labels” feature of iTOL. *De novo* elements in the clade were then classified as being part of the same family as the known element from RepBase (Figure S1), generating a TE library with many more elements having classifications.

This classified TE library as well as the full set of genomes were then used as input to RepeatMasker (Smit AFA, Hubley R, Green P 2013) version 4.1.1 with options ‘-gff -cutoff 200 -no_is -nolow -gccalc’ to generate copy number and positional data for TEs in all of the genomes. These hits were converted to fasta format using bedtools (Quinlan and Hall 2010) getfasta version 2.28.0 with the ‘-s’ option to force strandedness, then filtered once more for elements containing a TE coding sequence domain, as described previously. This produced TE annotations for each genome of elements that were predicted to be complete.

Data from the TE annotations on copy number of each TE family, total length each TE family occupies, and percentage of TE content were used for PCA, calculated using the prcomp (R Core Team) function and visualized with ggbiplot (Vu 2011) version 0.55 in R version 4.1.0. Point-Biserial correlation coefficients were calculated using the cor (R Core Team) function in R version 4.1.0 between binary (LineageGroup) and continuous (all other) variables from Table S1, and Spearman correlation coefficients were calculated between all pairs of continuous variables. For the binary LineageGroup variable, MoO and MoS genomes were assigned a value of 1, and MoT, MoL, and MoE genomes were assigned a value of 0.

Domain-based ML phylogenies of each TE family were constructed in the same way as those used to give *de novo* elements a family classification. The domains (with Pfam accession) used for each TE were: RVT_1 reverse transcriptase (PF00078.29) for *MAG_Ty3*, *Grasshopper*, *Ty3_MAG1*, *MGR583*, *PYRET*, and *MoTeR*, RVT_2 reverse transcriptase (PF07727.16) for *Copia_elem*, rve integrase (PF00665.28) for *MGRL3* and *Ty3_MAG2*, and DDE_1 transposase (PF03184.21) for *POT2* and *TcMar_elem*.

### Phylogeny of *M. oryzae* genomes

The genome tree of the *M. oryzae* isolates was generated by first annotating genes in each genome using FunGAP (Min et al. 2017) version 1.1.0 with arguments ‘--augustus_species magnaporthe_grisea --busco_dataset sordariomycetes_odb10’. RNAseq data for genome annotation was retrieved from the NCBI SRA database in June 2021. RNAseq for Guy11 (accession SRX5630771) was used as input for genomes of MoO and MoS lineage, RNAseq for B71 (accession SRX5900622) was used for MoT and MoL genomes, and RNAseq for MZ5-1-6 (accession SRX5092987) was used for MoE genomes. This resulted in predicted genes for each genome, which were input to OrthoFinder (Emms and Kelly 2019) version 2.5.4 along with the *M. grisea* NI907 proteome as the outgroup (retrieved from NCBI GenBank, accession GCA_004355905.1). OrthoFinder was run with options ‘-M msa -S diamond_ultra_sens -A mafft -T fasttree’, and the output identified 8,655 SCOs. These were aligned using MAFFT (Katoh and Standley 2013) version 7.312 with parameters ‘--maxiterate 1000 --globalpair’, and the alignments were concatenated. The ML phylogeny was produced from the alignment using RaxML (Stamatakis 2014) version 8.2.11 with options ‘-m PROTGAMMAGTR -T 24 -f a -x 12345 -p 12345 -# 100’, and was visualized in iTOL (Letunic and Bork 2019).

### Divergence analysis

To characterize RIP in *M. oryzae*, GC content was calculated using geecee from EMBOSS (Rice et al. 2000) version 6.6.0.0 in TEs and in coding sequences of the representative genomes. Median and median absolute deviation (MAD) values were calculated for each TE in each genome in R version 4.1.0 using the med and mad functions (R Core Team).

LTR divergence analysis was performed by first determining a consensus sequence for each flanking LTR. Elements from the *MAG_Ty3*, *Copia_elem*, *Grasshopper*, and *MGRL3* domain-based phylogenies were extracted from each representative genome, plus 1,000 bp on either side, using bedtools slop (Quinlan and Hall 2010) version 2.28.0. These sequences were then aligned using blastn from BLAST (NCBI Resource Coordinators et al. 2018) version 2.7.1+ against the clustered TE library from the intermediate step in the TE annotation pipeline, before LTRs were removed when filtering for domain containing elements (Figure S1). This helped to manually determine the element that best represented the LTR sequence of each TE, which was aligned, again using blastn (NCBI Resource Coordinators et al. 2018), back to the set of LTR-retrotransposon sequences plus flanking regions to extract LTRs. These extracted LTRs were then aligned using MAFFT (Katoh and Standley 2013) version 7.312, and a consensus sequence was generated using EMBOSS cons (Rice et al. 2000) version 6.6.0.0. The resulting LTR consensus sequences were used as the input library to RepeatMasker (Smit AFA, Hubley R, Green P 2013) version 4.1.1 with options ‘-gff -cutoff 200 -no_is -nolow -gccalc’, which produced positional information for all LTRs. This was used along with the original full sequence plus flanking regions to find which LTRs belonged to which full elements using bedtools intersect (Quinlan and Hall 2010) version 2.28.0. Finally, EMBOSS needle (Rice et al. 2000) version 6.6.0.0 was used to find the divergence of flanking LTR pairs.

Jukes-Cantor distance analysis was performed on all full-length TEs of interest, where the distance of each element to the consensus of its lineage, and to the consensus of all copies of that TE from any lineage were calculated. Following previous methods (Faino et al. 2016), we first produced the two types of consensus sequences by aligning TEs using MAFFT (Katoh and Standley 2013) version 7.312, then using EMBOSS cons (Rice et al. 2000) version 6.6.0.0 to generate the consensus of the alignment. The divergence of a TE from the consensus was found using EMBOSS needle (Rice et al. 2000) version 6.6.0.0, and this divergence was corrected by the Jukes-Cantor distance formula (Jukes and Cantor 1969). Using *Copia_elem* as an example, a consensus for all lineages was generated by aligning all copies of *Copia_elem* present in its domain-based ML phylogeny, then the distance of all *Copia_elem* from that consensus was found and plotted separately for each lineage. Also, a consensus was generated separately for *Copia_elem* from MoO, and the distance was computed as previously described, except using this consensus specific to the lineage. This was done for *Copia_elem* copies in each lineage separately and plotted. This process for making both plots was done for each of *MAG_Ty3*, *Grasshopper*, *POT2*, and *MGRL3* as well.

### Solo-LTR analysis

Solo-LTRs were identified by determining which LTRs (from the annotations previously generated for LTR divergence analysis) did not belong to an LTR-retrotransposon found by the TE annotation pipeline. Using the ‘-v’ option for bedtools intersect (Quinlan and Hall 2010) version 2.28.0 returned only the LTR sequences that had no overlap with an annotated TE, and thus were considered solo-LTRs. The number of solo-LTRs compared to the number of their full-length LTR-retrotransposon counterparts within and across genomes was used to determine whether the retrotransposon experienced expansion or removal from the genome.

### Analyses for investigating potential *POT2* HT

To investigate potential larger HT regions containing *POT2*, synteny analyses were performed between all *POT2* regions in Guy11 and B71. *POT2* sequences plus 50,000 bp on either side were extracted using bedtools slop and getfasta (Quinlan and Hall 2010) version 2.28.0. These regions were compared using nucmer and mummerplot from MUMmer (Marçais et al. 2018) version 4.0.0. To align the sequences, nucmer was called with the ‘--maxmatch’ option, and to visualize the alignment, mummerplot was called with options ‘--postscript --color’. This produced synteny plots that were visually screened through for long segments of synteny between Guy11 and B71 flanking the position of *POT2*.

In order to find any genes that may have been transferred along with *POT2*, gene trees produced by OrthoFinder based on amino acid sequence were screened to select those that follow the same topology as the *POT2* phylogeny. The ete2 (Huerta-Cepas et al. 2010) python package version 2.3.10 was used to determine which gene trees were structured such that the gene from Guy11 (MoO) and the gene from B71 (MoT) had the smallest distance from each other than from any other gene. Out of all SCOs, 388 genes had trees following this topology, and these were further refined by aligning their nucleotide sequences and determining topology in the same way as before. The remaining 38 genes whose trees based on nucleotide sequence followed this topology were visualized in IGV (Robinson et al. 2011) to determine any localization in the B71 genome.

### Investigating the potential region of recombination

The full segments of chromosome 7 from each representative genome that contained genes following a *POT2* topology were extracted using bedtools getfasta (Quinlan and Hall 2010) version 2.28.0, and the nucleotide sequences were aligned using MAFFT (Katoh and Standley 2013) version 7.312. A phylogeny was produced using RAxML (Stamatakis 2014) version 8.2.11, with options ‘-m GTRGAMMA -T 20 -f a -x 12345 -p 12345 -# 100’ based on the alignment.

To characterize the genes located in the potential region of recombination, we obtained their GO terms (Additional File 2) using the PANNZER (Törönen et al. 2018) webserver and their PFAM terms (Additional File 3) using pfam_scan.pl (Finn et al. 2016) version 1.6 with options ‘-e_dom 0.01 -e_seq 0.01’ against the Pfam-A.hmm library of HMMER (Eddy 2011) version 3.1b2. The output from PANNZER was then filtered for GO terms with PPV value > 0.6.

### Effector annotation and analysis

Effectors were predicted by following a previously established pipeline (Singh et al. 2019). Proteomes from FunGAP (Min et al. 2017) output were input into SignalP (Almagro Armenteros et al. 2019) version 5.0 to filter for proteins containing a signal peptide. The output of SignalP was then input to tmhmm (Krogh et al. 2001) version 2.0, which filtered out proteins containing a transmembrane domain. Finally, the remaining proteins were input to EffectorP (Sperschneider and Dodds 2022) version 3.0.

### Data processing and analysis

Analyses were conducted in a Linux environment with GNU bash version 4.2.46, GNU coreutils version 8.22, GNU Awk version 4.0.2, GNU grep version 2.20, and gzip version 1.5. Conda version 4.10.1 was used to install software. Scripts for parsing data were written in Python version 3.7.4, using biopython (Cock et al. 2009) version 1.79. R (R Core Team) version 4.1.0 was used to write scripts for data analysis and plotting, with packages ggplot2 (Wickham 2016) version 3.4.1, RColorBrewer (Neuwirth 2022) version 1.1-3, dplyr (Wickham H, François R, Henry L, Müller K, Vaughan D 2023) version 1.1.0, tidyverse (Wickham et al. 2019) version 2.0.0, and scales (Wickham H, Seidel D 2022) version 1.2.1.

### Availability of Data and Code

Datasets and intermediate analyses files are provided as additional data files, or available on Zenodo at DOI 10.5281/zenodo.7366416. Code and scripts used for all analyses are located in a GitHub repository (https://github.com/annenakamoto/moryzae_tes).

## Acknowledgements

We thank Dr. Anna Muszewska for advice on TE annotation, Dr. Pierre Gladieux for help with substitution rates in *M. oryzae*, and Dr. Michael Seidl for insight on using the Jukes-Cantor distance metric. We also thank Dr. Ursula Oggenfuss and Dr. Emile Gluck-Thaler for feedback on results. We thank members of the Krasileva Lab for feedback on manuscript preparation, and Dr. Pierre Gladieux, Dr. Ursula Oggenfuss, Dr. Emile Gluck Thaler, and Ivar Westerberg for helpful comments. This project utilized the Savio computational cluster resource provided by the Berkeley Research Computing program at the University of California, Berkeley.

## Contributions

AAN and PMJ developed the project with input from KVK. AAN designed methods, performed data collection and analyses, interpreted results, produced figures, and wrote the original manuscript draft, with guidance from PMJ. All authors reviewed, edited, and approved the final manuscript.

## Supplemental Material

Additional File 1: Supplementary Figures and Tables.

**Table S1:**
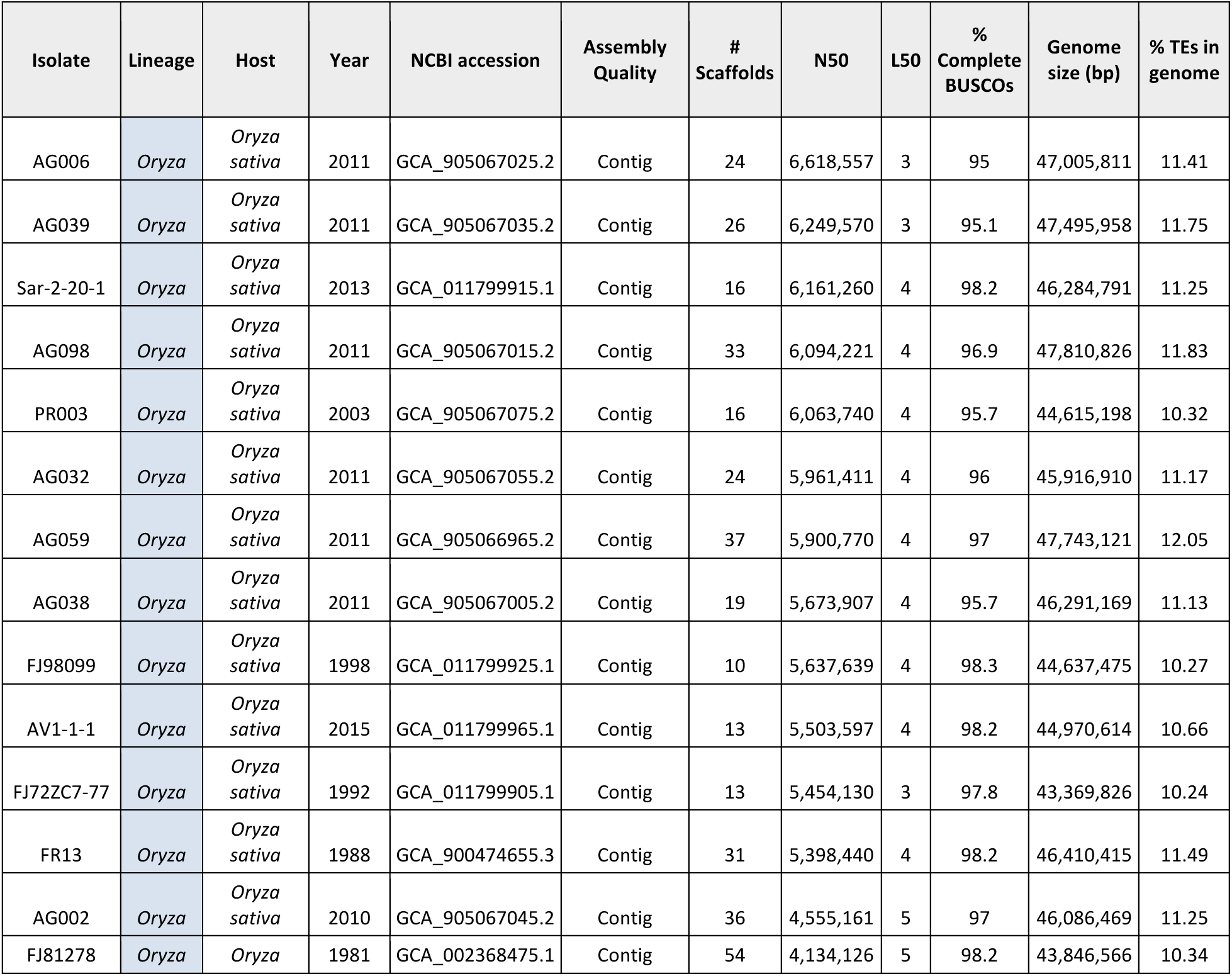
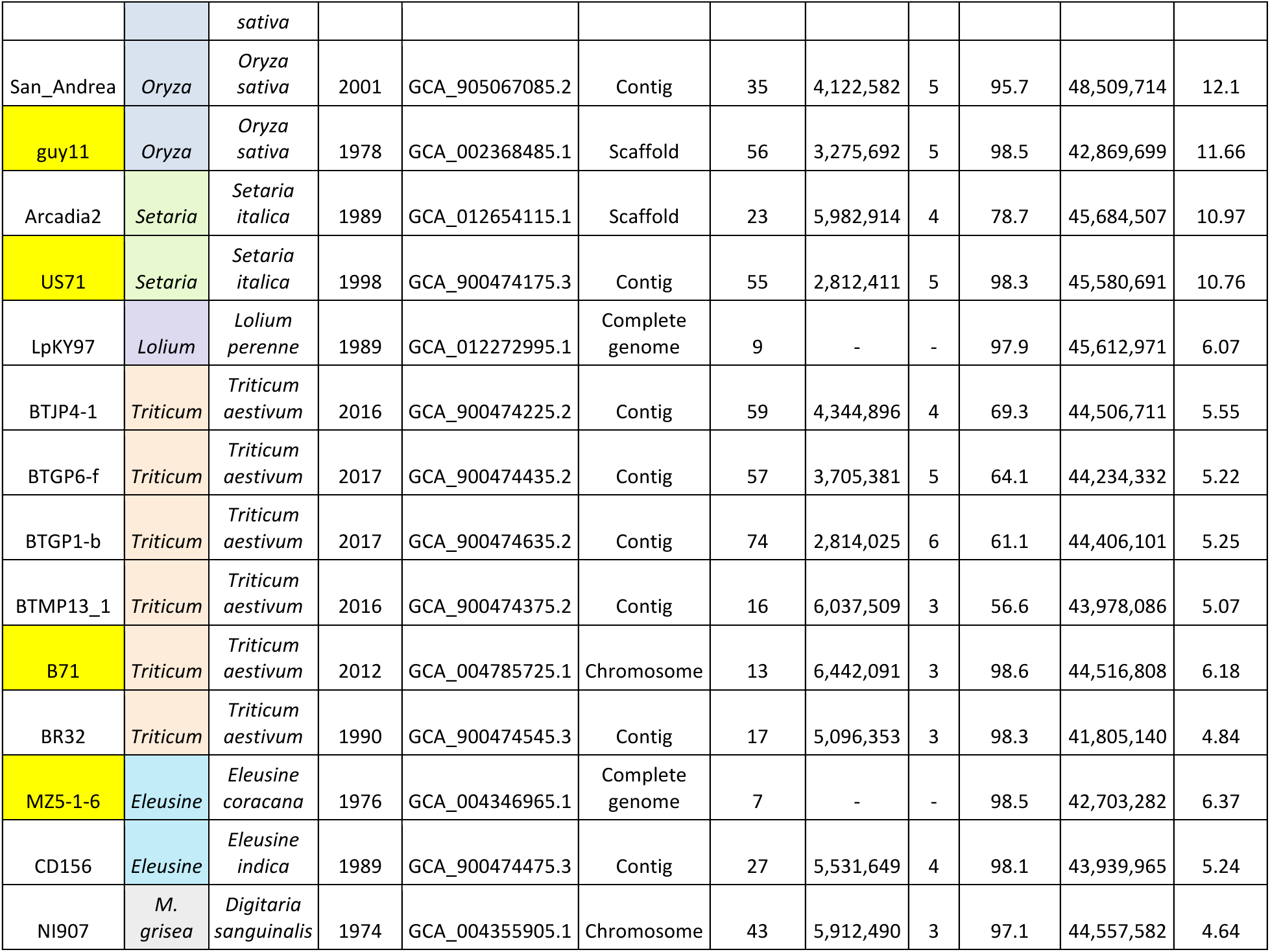
*M. oryzae* genomes used in this study, with the lineage they belong to, host they were isolated from, collection year, NCBI accession, assembly level, number of scaffolds, percentage of complete BUSCOs, genome size in base pairs, and the percentage of the genome containing TEs (as found by our TE annotation pipeline). The highest quality representative genome used for each lineage is highlighted in yellow. All genomes were retrieved from NCBI GenBank December 2020.

**Table S2:** Names and classifications of TEs discussed. The name we used throughout the paper for each TE family is shown, along with its original name in RepBase (Bao et al. 2015) fngrep version 25.10, the class, and the superfamily each element belongs to. We adopted a naming convention for *Ty3* (formerly *Gypsy*) elements, where any “*GY*” in the RepBase name was replaced with “*Ty3*” in order to use a non-discriminatory and respectful naming scheme (Wei et al. 2022). *Grasshopper* is the original name of the *GYPSY1* RepBase element, so it is used instead (Dobinson 1993). Elements that didn’t correspond to a specific family in RepBase are indicated by “N/A,” and are named by their superfamily (i.e. *Copia_elem*). LTR = long terminal repeat retrotransposon, NLTR = non-LTR retrotransposon, DNA = DNA transposon.

| TE family name used | RepBase name | Class | Superfamily |
| --- | --- | --- | --- |
| *Ty3_MAG1* | *GYMAG1* | LTR | *Ty3* |
| *Ty3_MAG2* | *GYMAG2* | LTR | *Ty3* |
| *Grasshopper (Grh)* | *Gypsy1* | LTR | *Ty3* |
| *MAG_Ty3* | *MAGGY* | LTR | *Ty3* |
| *MGRL3* | *MGRL3* | LTR | *Ty3* |
| *PYRET* | *PYRET* | LTR | *Ty3* |
| *Copia_elem* | N/A | LTR | *Ty1/Copia* |
| *MGR583* | *MGR583* | NLTR | *LINE/Tad1* |
| *MoTeR1* | *MoTeR1* | NLTR | *LINE/CRE* |
| *POT2* | *POT2* | DNA | *Tc1/Mariner* |
| *TcMar_elem* | N/A | DNA | *Tc1/Mariner* |

**Figure S1:**
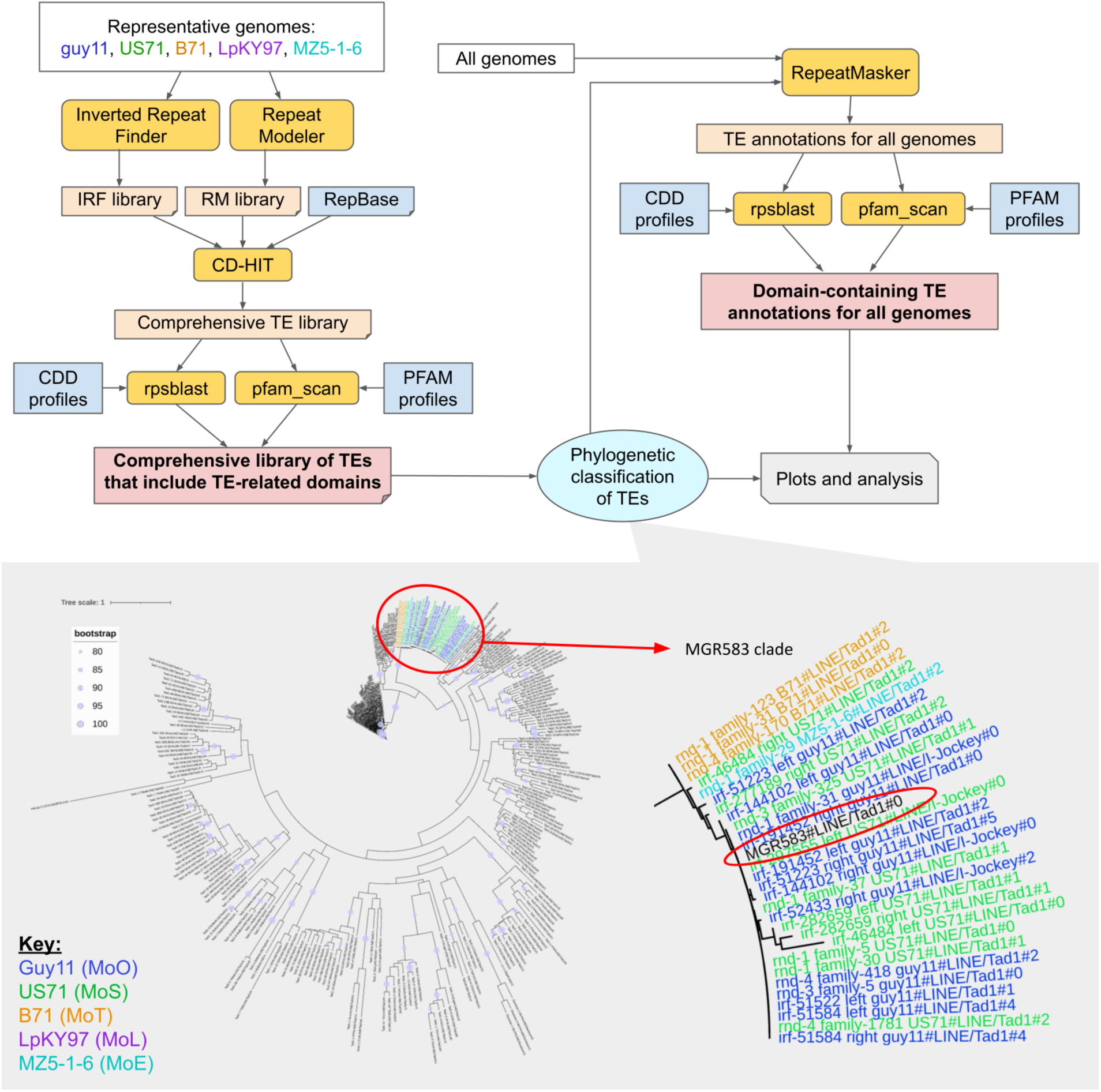
TE annotation pipeline diagram. The five representative genomes, Guy11 (MoO), US71 (MoS), B71 (MoT), LpKY97 (MoL), and MZ5-1-6 (MoE) were used for *de novo* TE annotation to produce an unbiased TE library that is representative of TE content in all lineages. The bottom gray box provides an example of how TEs were classified. Shown is a tree based on the Exo_endo_phos_2 domain with a phylogenetically defined TE subclade indicated by the red circle. Subclades of *de novo* elements (in color) that grouped with a known RepBase element (*MGR583* in this example, circled in red) were classified as that element’s family. Colored text names of *de novo* elements represent the genome they were annotated in, as shown in the key.

**Figure S2:**
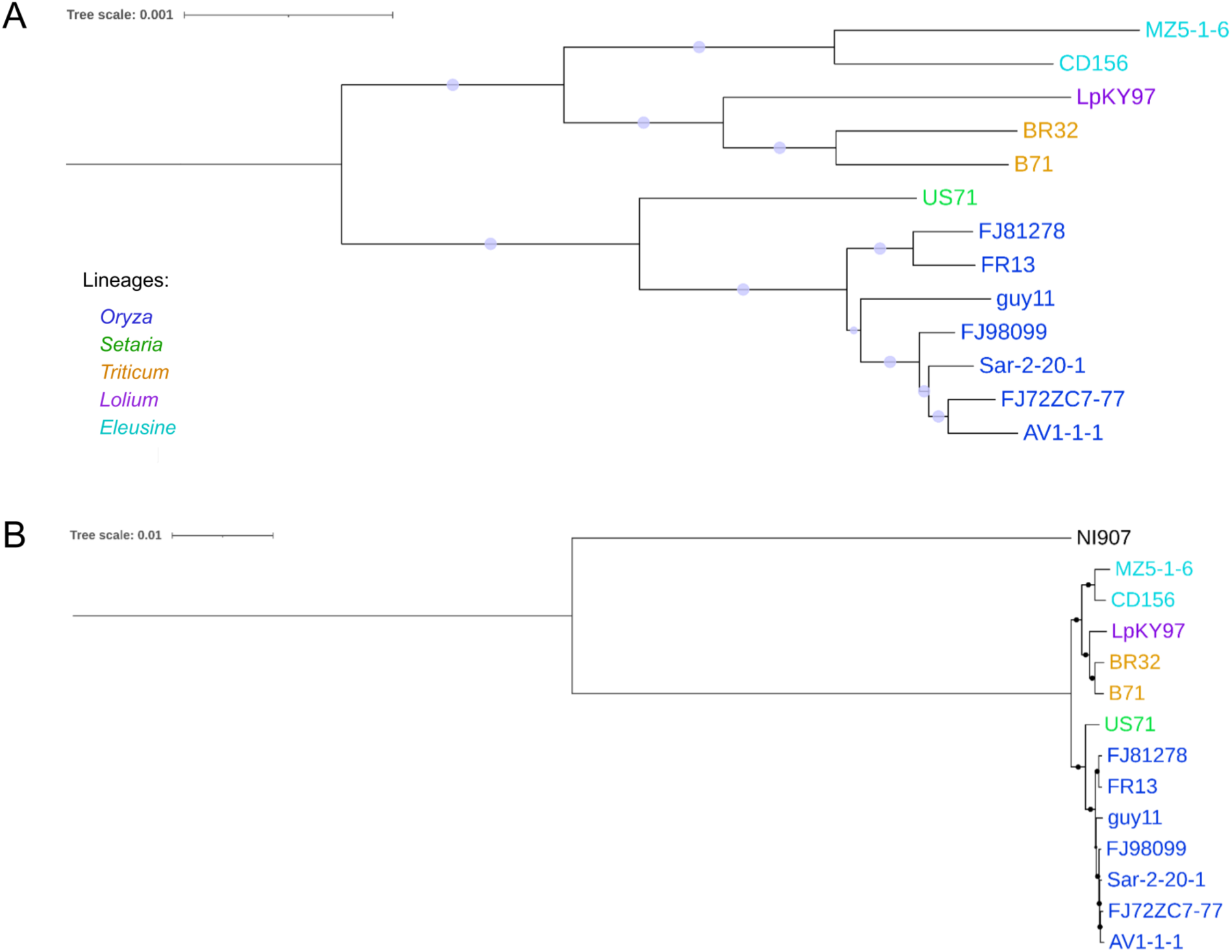
Maximum-likelihood (ML) phylogeny of *M. oryzae* genomes based on the alignment of 8,655 single copy orthologous genes (SCOs), **A,** zoomed in without outgroup and **B,** including *Magnaporthe grisea* outgroup (NI907). Only genomes with BUSCO score greater than 97% were included. Bootstrap value of 1 is indicated by circles on the branches.

**Figure S3:**
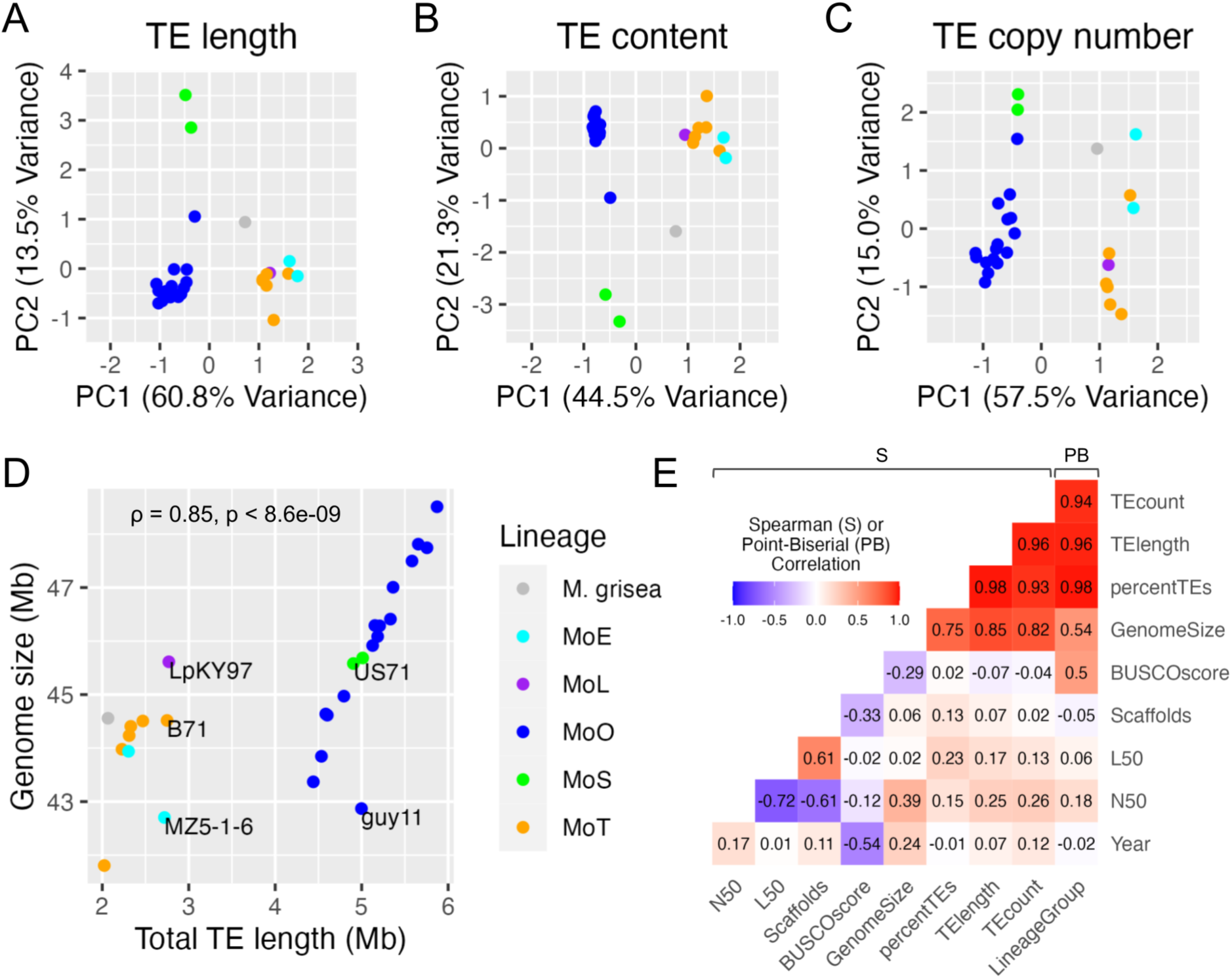
TE content clearly differentiates MoO-MoS versus MoT-MoL-MoE lineage groups, while other variables have weaker or no correlation to lineage identity. Scatterplots of principal components (PCs) 1 and 2 values from principal component analyses (PCAs) are shown, for **A,** the length that each TE occupies in each genome, **B,** the percentage of each TE’s copy number out of the total TE count in each genome, and **C,** the copy number of each TE in each genome. Each point represents one genome, and the percentage of variation that each PC describes is shown on the axes. **D,** The correlation between total TE length (Mb) and genome size (Mb) is shown, with Spearman’s ρ (rho) and p-value. **E,** Correlation matrix, where Point-Biserial correlation coefficients were calculated between binary (LineageGroup) and continuous (all other) variables, and Spearman correlation coefficients were calculated between all pairs of continuous variables.

**Figure S4:**
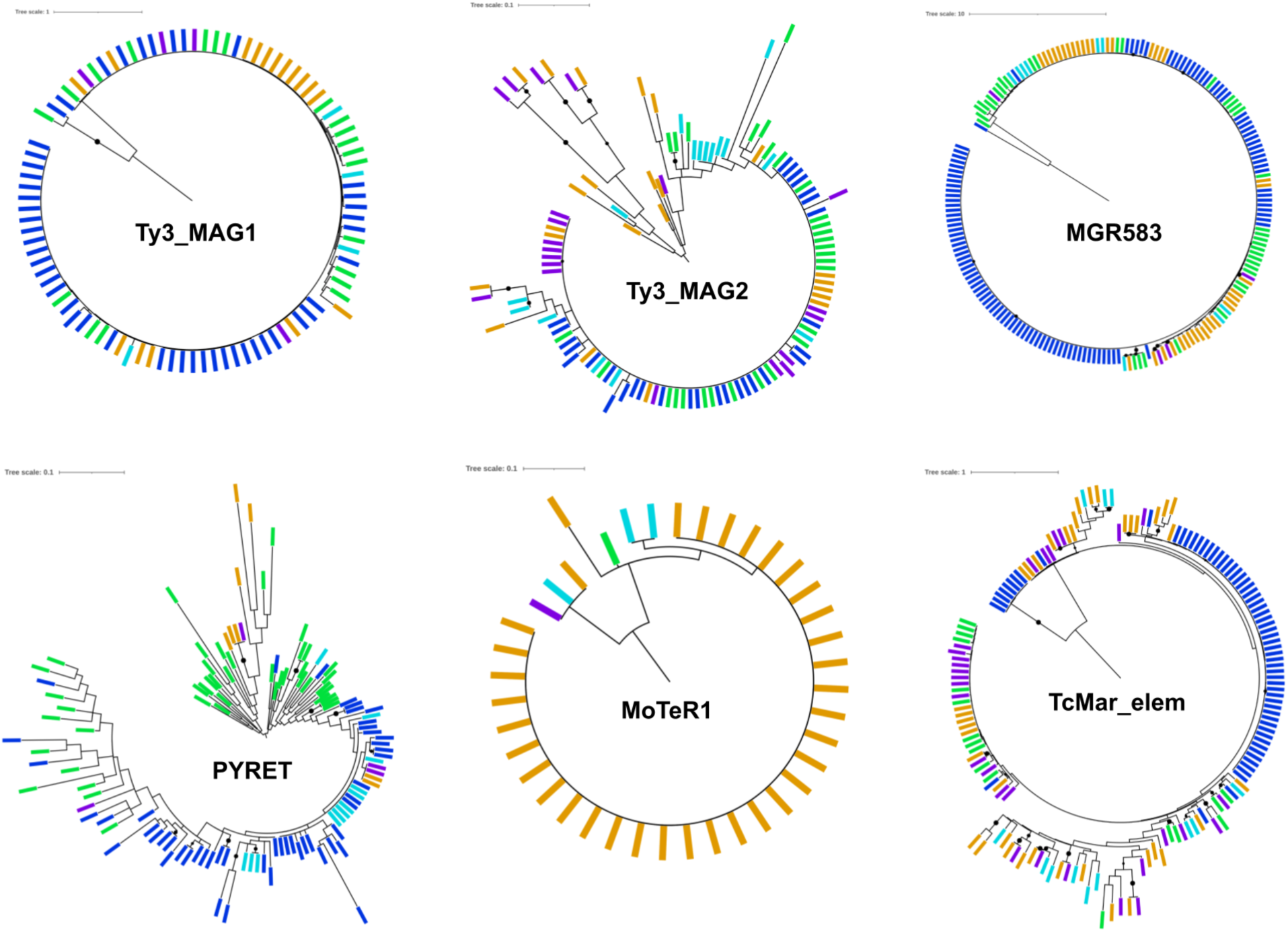
Domain-based maximum-likelihood (ML) phylogenies for *Ty3_MAG1*, *Ty3_MAG2*, *MGR583*, *PYRET*, *MoTeR1*, and *TcMar_elem*. Colored rectangle tips correspond to the genome each element is from: blue=Guy11 (MoO), green=US71 (MoS), orange=B71 (MoT), purple=LpKY97 (MoL), and cyan=MZ5-1-6 (MoE). Black circles indicate bootstrap value ≥80.

**Figure S5:**
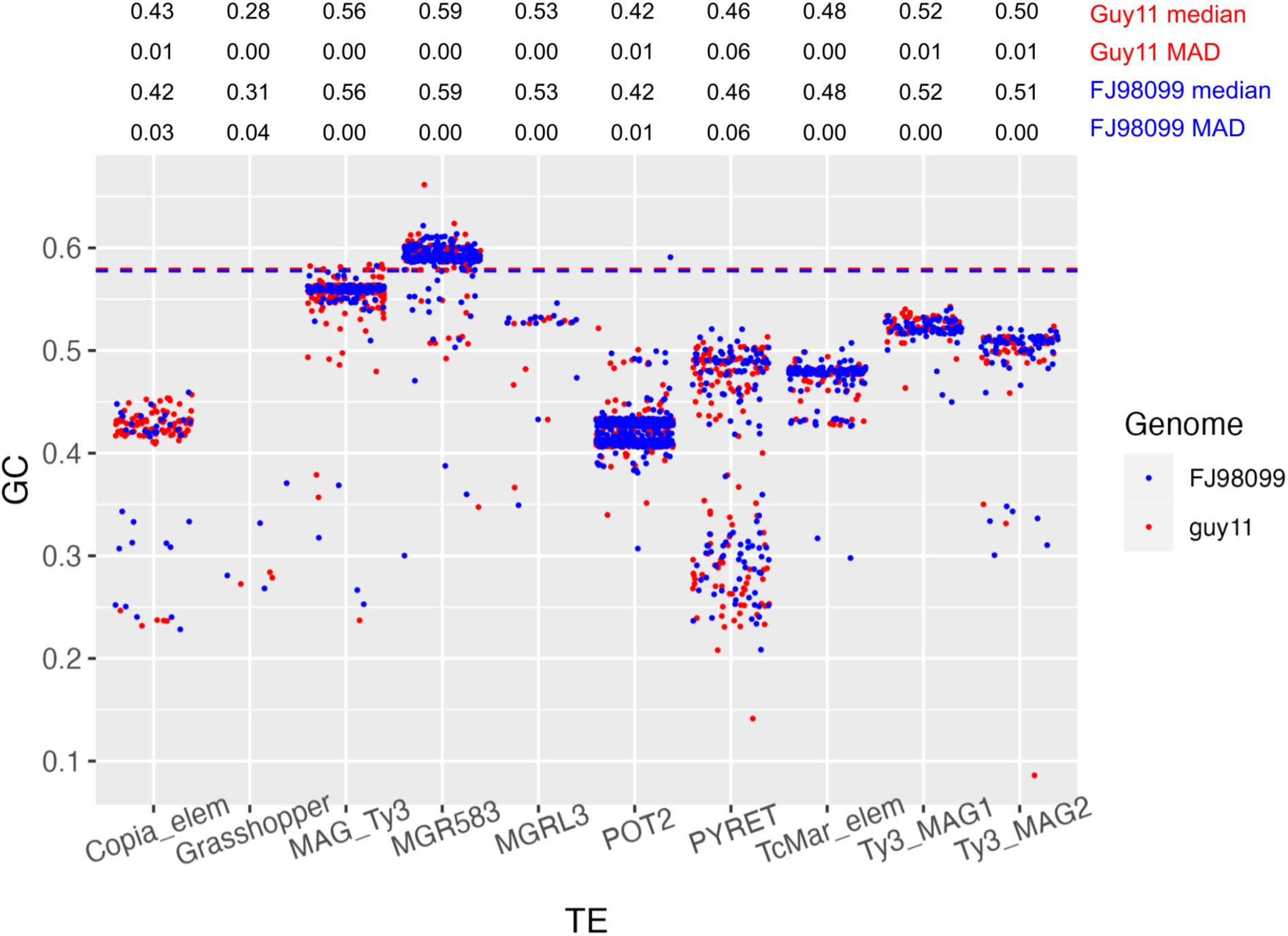
Jitter-plot showing GC content in each TE family, for the recombining Guy11 genome (red) and the clonal FJ98099 genome (blue). Each dot represents one TE copy, and dashed lines represent the genome-wide average GC-content of coding regions for each genome. Median GC content and median absolute deviation (MAD) values are displayed above for each TE in both genomes.

**Figure S6:**
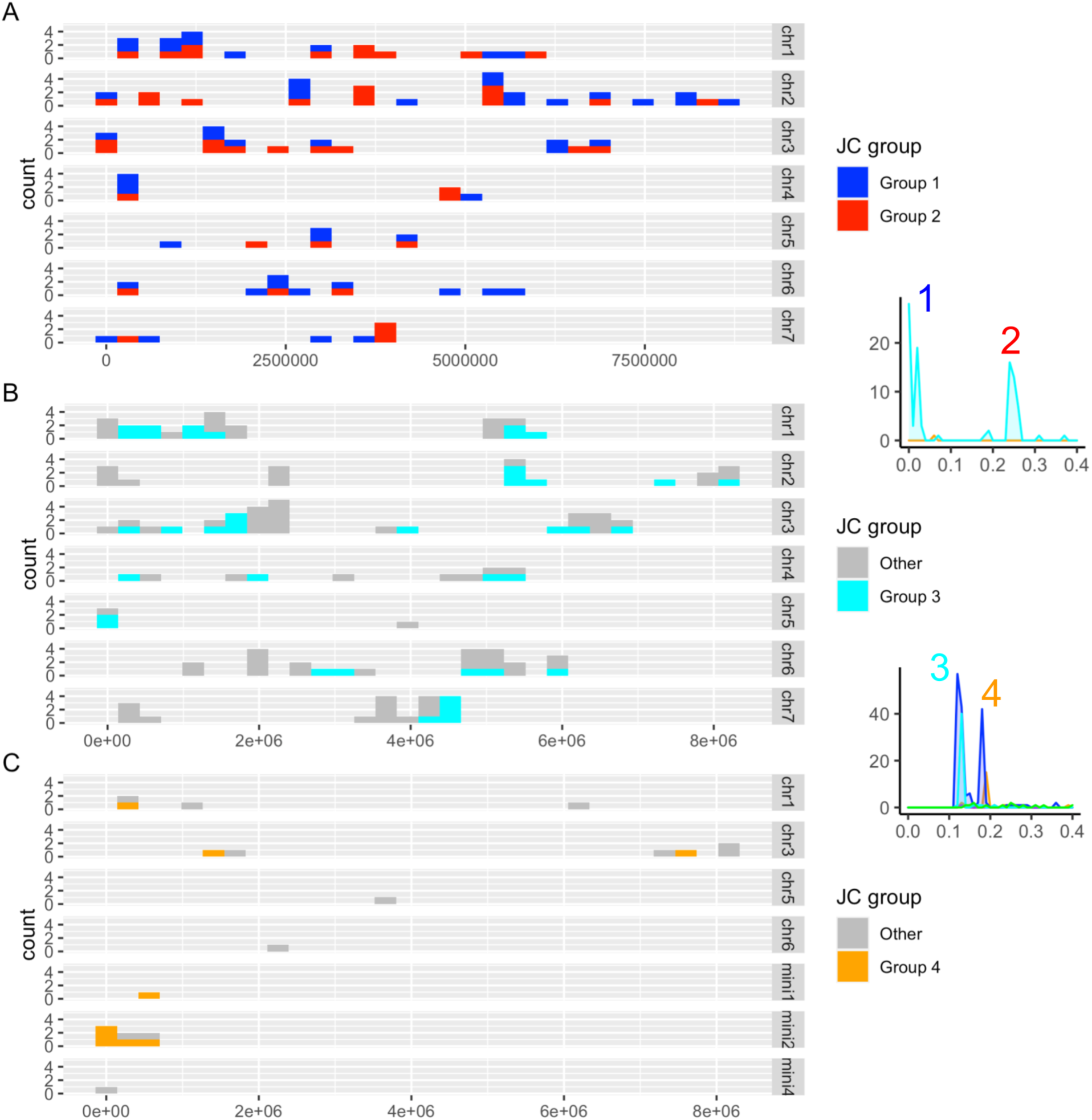
Localization of expanded TEs in *M. oryzae* genomes. **A,** The location of *Grasshopper* elements of lower (Group 1 in red) and higher (Group 2 in blue) Jukes-Cantor distance throughout MZ5-1-6’s seven chromosomes. The Jukes-Cantor plot for *Grasshopper* with groups 1 and 2 peaks labeled is shown for reference. **B,** The location of *POT2* elements in MZ5-1-6 that group with the lower Guy11 *POT2* Jukes-Cantor peak (Group 3 in cyan). **C,** The location of *POT2* elements in B71 that group with the higher Guy11 *POT2* Jukes-Cantor peak (Group 4 in orange). The Jukes-Cantor plot for *POT2* with groups 3 and 4 peaks labeled is shown for reference.

**Figure S7:**
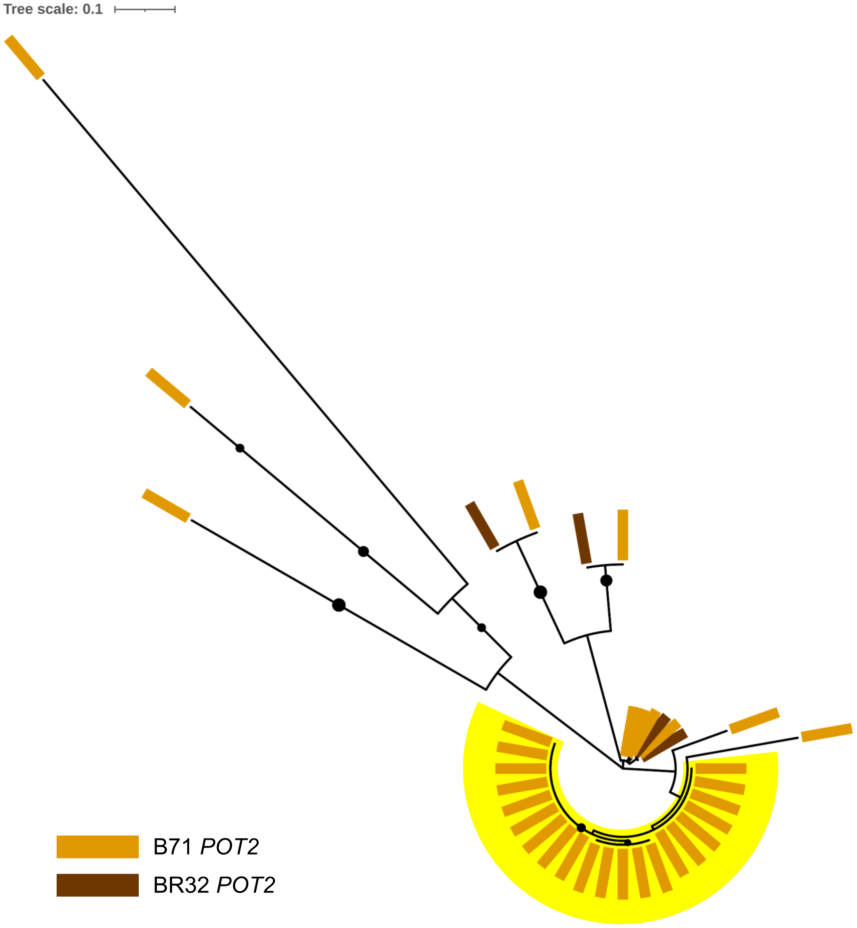
Domain-based maximum-likelihood (ML) phylogeny of *POT2* from MoT genomes B71 and BR32. The yellow highlighted clade corresponds to the potentially transferred B71 *POT2* clade from Figure 5A. Black circles indicate bootstrap values of ≥80.

**Figure S8:**
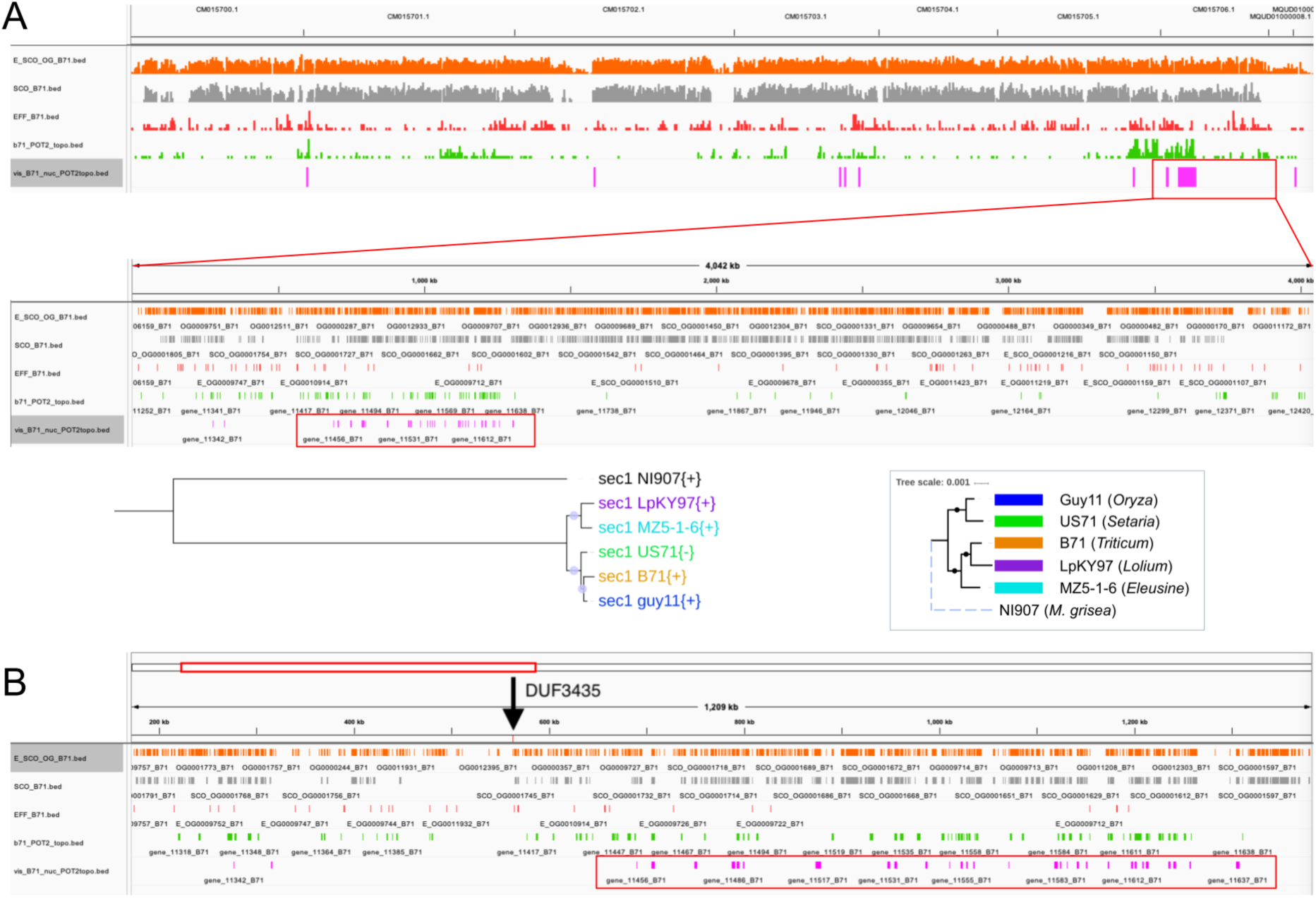
Genes following *POT2* tree topology are localized in a region on B71’s chromosome 7. **A,** The first IGV track shows all seven chromosomes, and the second track shows just chromosome 7. The tree on the left side shows the phylogeny constructed from an alignment of the full-length region in each isolate, and the genome tree is shown on the right for comparison. **B,** A gene containing a fragmented DUF3435 domain, which is associated with *Starship* elements, is located nearby and upstream of the region, indicated by the black arrow. The locations of genes that follow the *POT2* tree topology are indicated in pink, and the entire region is boxed in red in each track.

Additional File 2: GO terms output from PANNZER, filtered for >0.6 PPV value, for genes in the region on chromosome 7 following *POT2* topology.

Additional File 3: PFAM terms output from pfam_scan, filtered for E-value <0.01, for genes in the region on chromosome 7 following *POT2* topology.

